# Gain-of-function CCaMK in rice overrides genetic and anatomical barriers to arbuscular mycorrhizal colonisation

**DOI:** 10.64898/2026.05.21.726799

**Authors:** Gabriel Ferreras Garrucho, Raphaella Hull, Deven Rubens, Sarah Bowden, Ruth Bates, Matthew S. Hope, Emma Wallington, Uta Paszkowski

**Affiliations:** Crop Science Centre, Department of Plant Sciences, University of Cambridge, 93 Lawrence Weaver Road, Cambridge, CB3 0LE, UK; NIAB, 93 Lawrence Weaver Road, Cambridge, CB3 0LE, UK; RIKEN, Center for Sustainable Resource Science, Yokohama, Kanagawa 230-0045, Japan

**Author notes:** These authors contributed equally to this article (Gabriel Ferreras Garrucho, Raphaella Hull).

## Abstract

Arbuscular mycorrhizal (AM) symbiosis is conserved across land plants and is the default nutrient uptake strategy in nature. Within roots, AM colonisation is tightly patterned and dynamically tuned by nutritional cues. Multiple genetic modules contribute to this regulation, including the phosphate starvation response, DWARF14-LIKE (D14L) karrikin signalling, and the common symbiosis signalling pathway (CSSP). Transcriptional overlap among these has led to the hypothesis that phosphate starvation and D14L signalling act upstream of the CSSP. Here, we examined the epistatic relationship between D14L and CSSP in rice. Overexpression of an autoactive gain-of-function CCaMK (*gofCCaMKox*) restored AM colonisation and symbiosis marker gene expression in *d14l* mutants to wild-type levels or above, whereas overexpression of wild-type CCaMK did not, confirming that CSSP operates downstream of D14L signalling. However, *gofCCaMKox* did not rescue the *d14l* mesocotyl elongation phenotype, supporting a bifurcation of D14L into developmental and symbiotic outputs. Unexpectedly, *gofCCaMKox* also expanded fungal access to normally restrictive tissue domains (the meristematic zone and endodermis) assigning a role for CCaMK activation in defining root zone and cell-type competence for AM colonisation. Despite restored colonisation, introduction of *gofCCaMKox* into *d14l* produced arbuscules, which however were less developed and had increased hyphal septation, revealing a CCaMK-independent role for D14L in intraradical colonisation and arbuscule development. Transcriptome profiling resolved AM-relevant genes into modules controlled by CCaMK activation alone, in combination with D14L, or requiring additional colonisation-associated cues, and further suggested CCaMK primarily acts through AP2 transcription factors. Together, these findings reinforce CCaMK as a master regulator of AM symbiosis at the genetic, transcriptomic and anatomical levels while uncovering CCaMK-independent functions of D14L in arbuscule development.

## Introduction

Arbuscular mycorrhizal (AM) symbiosis is an ancient, widespread mutualism between land plants and fungi of the Glomeromycotina. In this interaction, AM fungi enter and expand within plant roots, leading to the formation of arbuscules, highly branched hyphal structures within cortical cells, where bidirectional nutrient exchange occurs: plants acquire mineral nutrients (notably phosphate) in return for photosynthates (sugars and lipids), facilitated by the large membrane interface at arbuscules (Smith and Read, 2008).

Due to this intimate nature and nutritional importance, AM symbiosis is tightly controlled by the plant. Spatially, AMF enter through epidermis and exodermis, proliferate through the cortex, most commonly via the apoplast, and exclusively invade cortical cells to form arbuscules (Smith and Read, 2008). The endodermis is not known to support AM fungal colonisation, and the fungus never invades the vascular cylinder; colonisation is also absent the meristematic region of the root tip and aerial tissues in angiosperms (Alexander et al., 1988; Parniske, 2008; Smith and Read, 2008; Genre et al., 2020; Orozco-Mosqueda et al., 2026). The mechanisms defining these permissive and restrictive domains remain largely unknown. A recent study shed some light on the mechanism behind the inner cortical cell development and identity in *Medicago truncatula*, where arbuscules are exclusively hosted. DELLA proteins, mobile from endodermis-stele to cortex, were shown to regulate the number of these arbuscule-hosting inner cortical cells (An et al., 2025). However, the reason behind the inability of the outer cortex to form arbuscules remains a mystery (Floss et al., 2013; An et al., 2025).

AM symbiosis is also dynamically regulated according to host status and environment, particularly nutrient availability (Kiers et al., 2011; Zheng et al., 2015; Kobae et al., 2016; Wang et al., 2017; Whiteside et al., 2019). Under high phosphate, colonisation is strongly repressed (Mosse et al., 1973; Koide and Mosse, 2004), a response linked to the inhibition of the phosphate starvation response (PSR) and its master regulators PHOSPHATE STARVATION RESPONSE (PHR) (Shi et al., 2021; Das et al., 2022; Paries and Gutjahr, 2023; Zhang et al., 2024). A second major regulatory input is D14L/KAI2 signalling (reviewed in Hull et al., 2021). The α/β-fold hydrolase Dwarf14-Like (D14L) is the rice homologue of KArrikin Insensitive (KAI2), the karrikin (KAR) receptor in *Arabidopsis thaliana* (Kagiyama et al., 2013), and the older paralogue of the SL receptor Dwarf14 (D14)(Delaux et al., 2012). D14L is essential for symbiosis in rice: *d14l* mutants fail to mount AM-associated transcriptional responses and do not support intraradical colonisation (Gutjahr et al., 2015), with related phenotypes reported across angiosperms (Liu et al., 2019; Li et al., 2022; Meng et al., 2022). D14L signals via the SCF component DWARF3 (D3) to promote degradation of the repressor SMAX1 (Nelson et al., 2011; Yoshida et al., 2012; Stanga et al., 2013; Stanga et al., 2016; Choi et al., 2020; Khosla et al., 2020; Zheng et al., 2020). In rice, SMAX1-regulated transcriptomes implicate D14L signalling in priming multiple symbiosis-relevant processes, including strigolactone biosynthesis (*D17/CCD7* and *D10/CDD8*), perception of microbial signals (chitin oligomer co-receptor *NFR5*), and components of the common symbiosis signalling pathway (CSSP) such as *SYMRK*, *CCaMK*, and *CYCLOPS* (Choi et al., 2020; Hull et al., 2021). Consistently, symbiotic Ca^2+^-spiking in response to CO4 and LCOs was shown to be affected in the *d14l* mutant, further suggesting that D14L signalling controls the activation of the CSSP (Li et al., 2022).

At the centre of the CSSP sits CCaMK, which decodes symbiotic Ca^2+^ oscillations and activates downstream transcriptional programs via phosphorylation of CYCLOPS (Ané et al., 2004; Mitra et al., 2004; Yano et al., 2008; Swainsbury et al., 2012; Miller et al., 2013; Singh et al., 2014). Gain-of-function (gof) CCaMK variants, containing only the kinase domain or mutations of the autophosphorylated threonine residue, can bypass upstream CSSP defects in legumes, restoring symbiosis and inducing AM marker genes where wild-type CCaMK cannot (Gleason et al., 2006; Tirichine et al., 2006; Yano et al., 2008; Hayashi et al., 2010; Madsen et al., 2010; Miller et al., 2013; Takeda et al., 2015). These observations lend support for CCaMK activation as a critical checkpoint for symbiosis establishment.

Here, we sought to assess the epistatic relationship between D14L and the CSSP for regulation of AM symbiosis in rice. To this end, we overexpressed native CCaMK or an autoactive version (*gofCCaMKox*) in the *d14l* mutant, allowing us to determine whether *CCaMK* expression alone, or its further activation by Ca^2+^-spiking, sit downstream of D14L signalling. AM colonisation was fully restored to wild-type levels only by *gofCCaMKox*, while overexpression of native *CCaMK* failed to rescue intraradical colonisation in the *d14l* mutant. This indicates that CCaMK activation, not just *CCaMK* expression, is compromised in *d14l* and places CSSP activation downstream of D14L signalling in root permissiveness to AM fungi. Despite recovery of intraradical colonisation, complemented lines displayed smaller arbuscules and increased hyphal septation, suggesting additional D14L functions in intraradical growth and arbuscule development that are CCaMK-independent. Strikingly, *gofCCaMKox* also enabled colonisation beyond canonical permissive domains, including arbuscules in endodermal cells and extensive colonisation of the meristematic zone of the root tip. Finally, transcriptome profiling distinguished gene sets controlled by CCaMK activation versus D14L signalling independently of the CSSP, including nutrient transporters such as *STR2* and *PT11*, providing a molecular framework for these phenotypes.

## Results

### CCaMK activation is downstream of D14L signalling for AM symbiosis

The epistatic relationship between D14L signalling and CCaMK was investigated. Based on the upregulation of *CCaMK* expression in rice *smax1* (Choi et al., 2020), we examined whether reduced *CCaMK* expression in the *d14l* mutant could explain the loss of AM symbiosis. To this end, overexpression constructs of native *CCaMK* were generated in rice in wild-type (WT), *ccamk*, and *d14l* backgrounds (WT*^CCaMKox^*, *ccamk^CCaMKox^*, and *d14l^CCaMKox^*, respectively) (Fig. 1A, table S1, S2). The WT and *ccamk* complementation constructs served as positive controls to assess the functionality of the constructs and to identify potential side effects of *CCaMK* overexpression. Overexpression lines of *eGFP* under the same promoter in wild-type (WT) and *d14l* backgrounds were used as a tissue culture, transformation, and transgene expression controls (WT*^GFP^*, *d14l^GFP^*).

**Figure 1.**
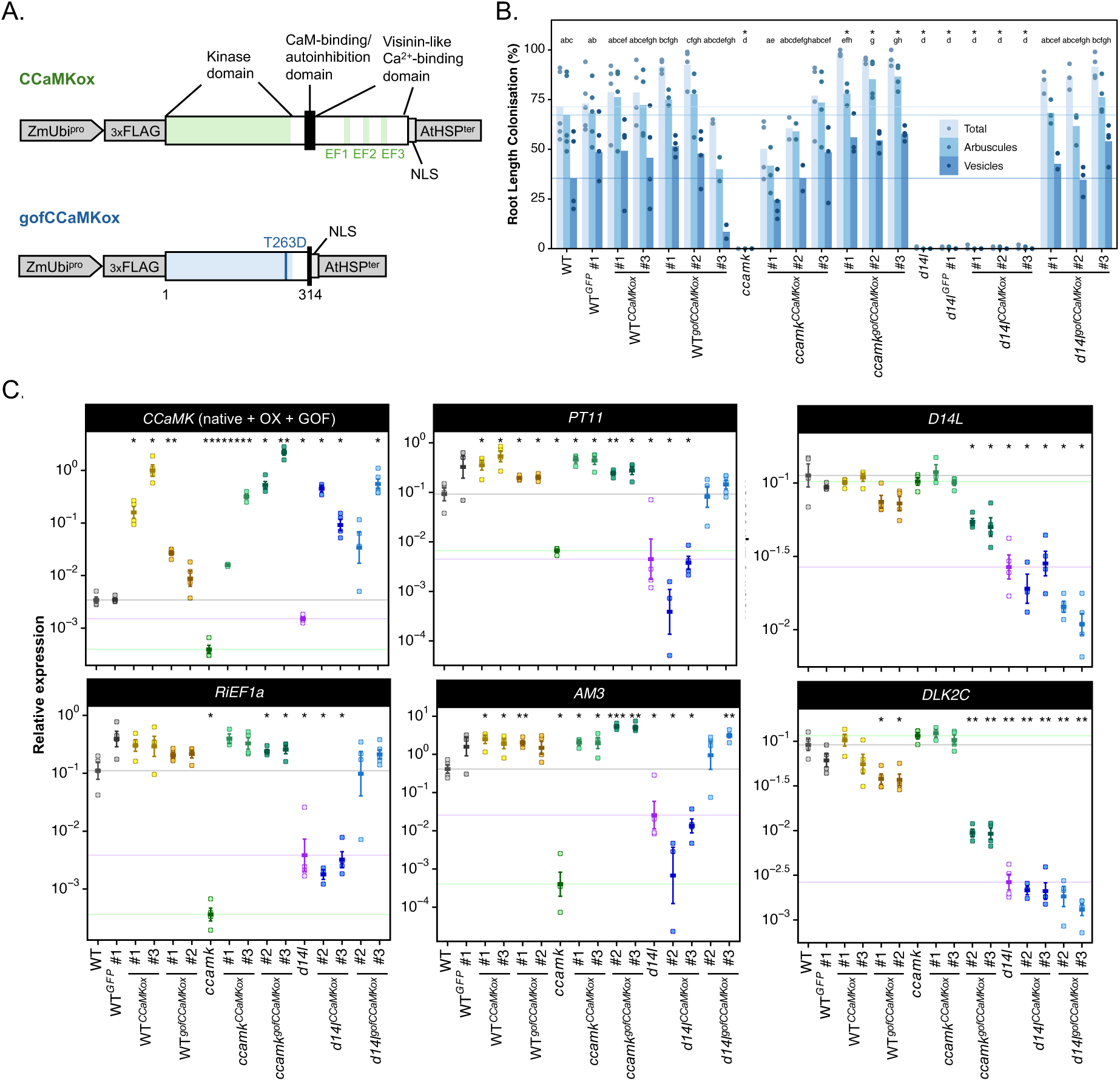
*gofCCaMKox*, but not *CCaMKox*, restores AM colonisation and symbiotic gene expression in *d14l*. (A) CCaMK overexpression constructs used for rice transformation in this study. For the *CCaMKox* construct, native CCaMK consisting of kinase domain (residues 13-298), calmodulin (CaM)-binding domain (321-334), coiled coil domain (343-363), EF-hand 1 (392-427), EF-hand 2 (428-463), and EF-hand 3 (470-505) was expressed under the promoter of *Zea mays* Ubiquitin (*ZmUbi*) promoter, with an N-terminal *3xFLAG* tag, a C-terminal nuclear localisation signal (NLS, from SV40 virus, PKKKRKV, (Lu et al., 2021)) and with the terminator from *AtHSP*. The *gofCCaMKox* construct generated in this study consisted of a protein truncation (1-314) and a phosphomimetic replacement of the autophosphorylation threonine by aspartic acid (T263D), under the same promoter, tags and terminator as *CCaMKox*. (B) Quantification of AM colonisation for *CCaMKox* and *gofCCaMKox* lines colonised by *R. irregularis* at 7 weeks-post-inoculation (wpi). Individual data points displayed, bars represent means for each genotype and structure. Horizontal lines show the mean for total colonisation, arbuscules, and vesicles for WT. Statistically significant differences are determined by Kruskal-Wallis test followed by Pairwise Wilcoxon rank sum test. Letters denote statistically significant differences between groups (*p* < 0.05), asterisks indicate significant differences to WT (*, *p* < 0.05; **, *p* < 0.01; ***, *p* < 0.001; ****, *p* < 0.0001), regarding total colonisation. (C) Normalised gene expression levels determined via qRT-PCR for *CCaMKox* and *gofCCaMKox* lines at 7 wpi. Each graph represents gene expression of a particular gene or product defined by a primer pair (table S2), gene name or product on top, normalised to the geometric mean of three housekeeping genes. Individual datapoints are shown, error bars represent mean ± standard error. Point colour represents genotype. Horizontal lines show the mean for WT (grey), *ccamk* (green) and *d14l* (purple). Statistically significant differences are determined by Anova, followed by two-sided t-test (*, *p* < 0.05; **, *p* < 0.01; ***, *p* < 0.001; ****, *p* < 0.0001), only comparisons with WT are shown.

The colonisation phenotypes at 7 weeks-post-inoculation (wpi) of 2-3 independent transgenic lines of WT*^CCaMKox^*, *ccamk^CCaMKox^*, and *d14l^CCaMKox^*were compared against their respective background genotypes (Fig. 1B, S1, repeated in an independent experiment in Fig. S2A, table S3). WT total colonisation was 71.40% ±7.70% and WT*^GFP^* total colonisation was 73.00% ±6.16%. Total colonisation was similar in all three WT*^CCaMKox^* lines and not statistically different to WT or WT*^GFP^*, averaging 78.63% ±5.43%. No intraradical colonisation was observed in *ccamk*, reproducing the phenotype reported in rice (Gutjahr et al., 2008). WT total colonisation was restored in three independent *ccamk^CCaMKox^* lines (*p <* 0.05), with an average of 63.00% ±5.78%, demonstrating that the transgenic *CCaMK* construct was fully functional. The variation in complementation observed for *ccamk^CCaMKox^* #1-3 positively correlated with the level of transgene expression (Fig. 1C, S2B, table S4). In contrast, *CCaMKox* was unable to rescue intraradical colonisation in *d14l* in three independent lines, whereby no significant difference in colonisation was observed between *d14l^CCaMKox^* and *d14l* (*p* > 0.05), all lines having <1% total colonisation, that is, extremely rare colonisation events, as previously reported for *d14l* mutants (Choi et al., 2020). The levels of *CCaMKox* expression in *d14l* were comparable to the WT and *ccamk* background (Fig. 1C, S3, table S4), where, in both cases, WT levels of colonisation were observed. Gene expression quantification by RT-qPCR for known plant marker genes induced during intraradical colonisation by AMF, *PT11*, *AM3*, *AM1* (Gutjahr et al., 2008) as well as the *Rhizophagus irregularis* constitutive gene *RiEF1a*, confirm the phenotypic results. *CCaMKox* expression allowed restoration of marker gene expression to levels statistically equivalent or superior to the WT in *ccamk*, but not in *d14l* (Fig. 1C, independent experiment in Fig. S2B, table S4). Together, these results suggest that D14L-mediated upregulation of *CCaMK* expression cannot solely explain D14L function in symbiosis.

Based on these results, the up-regulation of chitin oligomer receptor *NFR5* and further CSSP components such as *SYMRK* in *smax1* (Choi et al., 2020), as well as Ca^2+^-spiking being reduced in the *d14l* mutant (Li et al., 2022), we hypothesised that CCaMK activation, not only expression, is compromised in *d14l*. Thus, we tested whether constitutive activation of CCaMK could restore symbiosis in rice *d14l*. A gain-of function CCaMK (*gofCCaMK*) construct was designed, consisting of a protein truncation (1-314) and replacement of the autophosphorylation threonine by aspartic acid (T263D)(Fig. 1A, table S1, S2), shown in previous studies to auto-activate CCaMK function in legumes leading to spontaneous nodule formation and expression of AM marker genes (Gleason et al., 2006; Tirichine et al., 2006; Yano et al., 2008; Hayashi et al., 2010; Madsen et al., 2010; Miller et al., 2013; Takeda et al., 2015). This construct was overexpressed in WT, *ccamk*, and *d14l* backgrounds.

Total AM intraradical colonisation at 7 wpi of WT*^gofCCaMKox^*lines averaged at 86.50% ±4.16%, higher than WT and WT*^GFP^*, although not statistically significant (*p* > 0.05). Similarly, total and arbuscule colonisation in *ccamk^gofCCaMKox^* lines were significantly higher than in *ccamk* and restored to levels higher than WT and *ccamk^CCaMKox^*, but only statistically significant for comparisons with WT, with an average of 96.09% ±0.77%. The full complementation of symbiosis observed in *ccamk^gofCCaMKox^* lines demonstrated that the transgenic construct was functional for AM symbiosis. In contrast to the failure of *CCaMKox* to rescue symbiosis in *d14l*, *gofCCaMKox* restored colonisation in *d14l*, including arbuscule development, in three independent lines, with an average total colonisation of 87.70% ±2.01%, and 69.5% ±3.02% of arbuscules (Fig. 1B, S1, independent experiment in Fig. S2A, table S3).

Gene expression quantification by RT-qPCR for known plant marker genes induced during intraradical colonisation by AMF, *AM1, AM3*, and *PT11*, as well as the *R. irregularis* housekeeping gene *RiEF1a*, confirm the microscopic phenotyping results. *gofCCaMKox* lines reached marker gene expression levels statistically equivalent or superior in all backgrounds, including *d14l* (Fig. 1C, independent experiment in Fig. S2B). Conversely, expression levels of *D14L*, as well as the reporter genes for D14L signalling activity, *DLK2A* and *DLK2C*, were not affected by expression of *gofCCaMKox* (Fig. 1C, S2, S3). This indicates that D14L signalling is not retroactively regulated by the CSSP, and that D14L regulates genetic components independently of CCaMK, such as *DLK2s*. Interestingly, *NSP2* was down-regulated in the *d14l* but not in the *ccamk* mutant, and was activated by *gofCCaMKox* but not by *CCaMKox*, thus suggesting regulation both by D14L and active CCaMK, independently of colonisation levels (Fig. S3).

**Figure 2.**
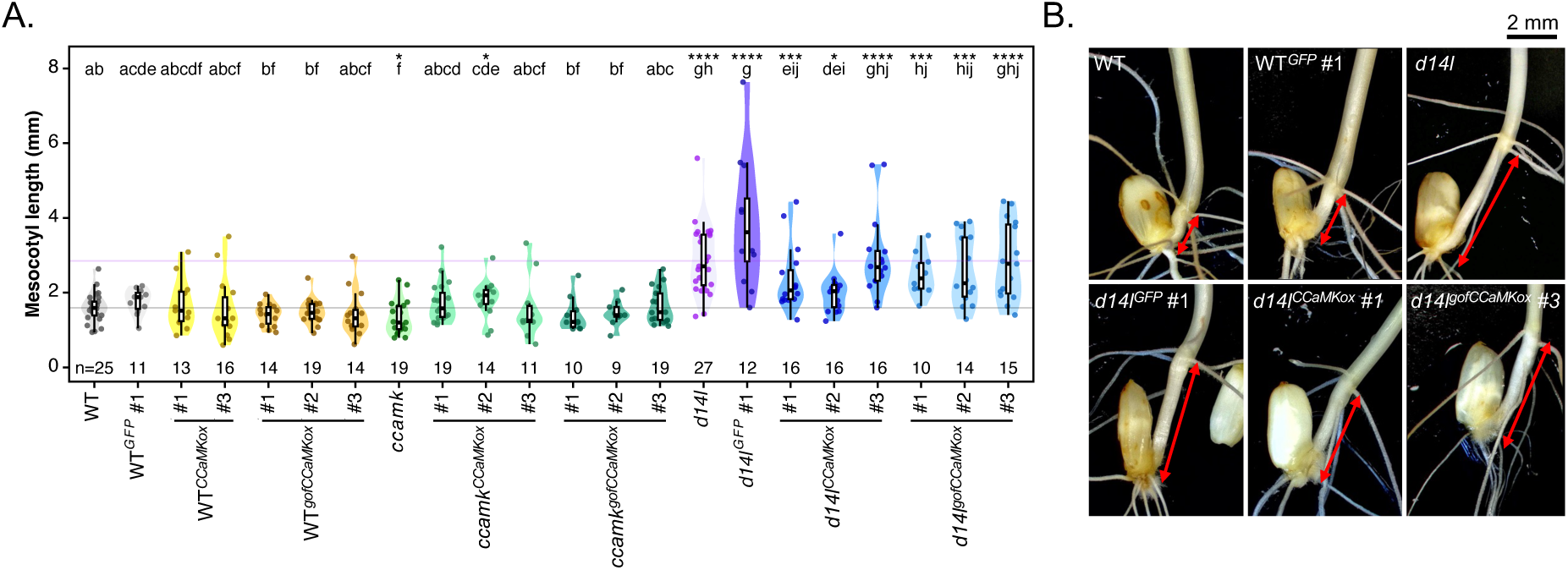
D14L control of mesocotyl elongation is not rescued by CCaMK overexpression or autoactivation. (A) Quantification of mesocotyl length for *CCaMKox* and *gofCCaMKox* lines at 7 days-post-imbedding (dpi). Individual datapoints are shown (n > 8 seedlings, indicated on the graph). Point colour represents genotype. Horizontal lines show the mean for WT (grey) and *d14l* (purple). Statistically significant differences are determined by Kruskal-Wallis test, *p*-value displayed, followed by Pairwise Wilcoxon rank sum test. Letters denote statistically significant differences between groups (*p* < 0.05), asterisks indicate significant differences to WT (*, *p* < 0.05; **, *p* < 0.01; ***, *p* < 0.001; ****, *p* < 0.0001). (B) Representative stereomicroscopy images of seedlings at 7 dpi, mesocotyl marked by red arrows.

**Figure 3.**
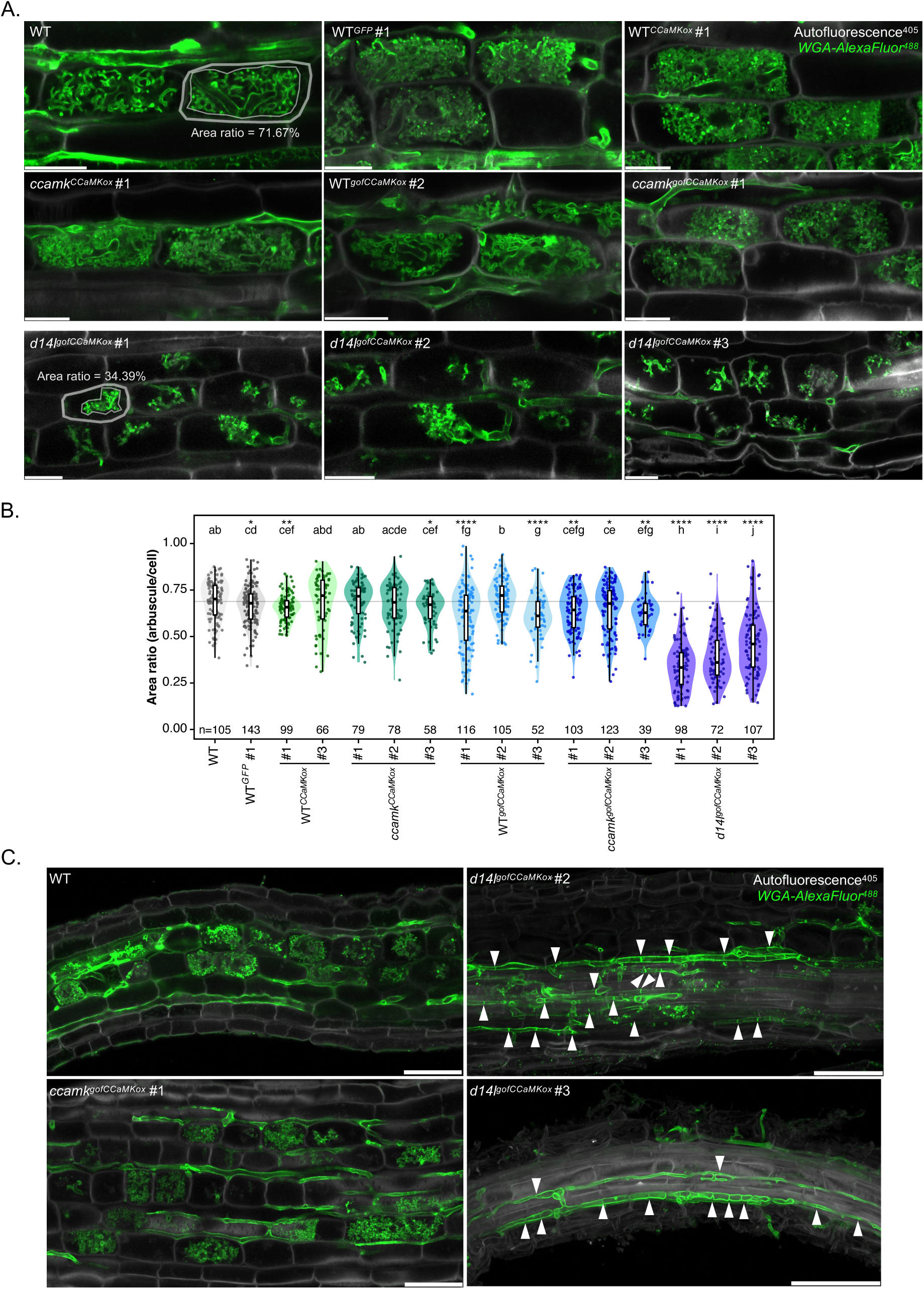
*d14l^gofCCaMKox^* roots show reduced arbuscule size and increased septation of intraradical hyphae. (A) CSLM of WGA-AlexaFluor^488^ stained roots of *CCaMKox* and *gofCCaMKox* lines colonised by *R. irregularis* at 7 weeks-post-inoculation (wpi) showing smaller, less-branched arbuscules exclusively in *d14l^gofCCaMKox^* lines. At least two independent transformant lines, three plants, three roots, and three arbuscules of each genotype were imaged. One representative image for each genotype is shown, one plane corresponding to the point of maximum arbuscule expansion. White channel corresponds to autofluorescence after excitation at 405 nm, green to WGA-AlexaFluor^488^. Scale bar, 20 μm. (B) Quantification of arbuscule size distribution of *CCaMKox* and *gofCCaMKox* lines colonised by *R. irregularis* at 7 wpi. Individual datapoints are shown (*n* > 50 arbuscules, specified on the graph, from ≥ 3 biological replicates). Point colour represents genotype. Horizontal lines show the mean for WT (grey) and *d14l* (purple). Statistically significant differences are determined by Kruskal-Wallis test, *p*-value displayed, followed by Pairwise Wilcoxon rank sum test. Letters denote statistically significant differences between groups (*p* < 0.05), asterisks indicate significant differences to WT (*, *p* < 0.05; **, *p* < 0.01; ***, *p* < 0.001; ****, *p* < 0.0001). (C) CSLM of WGA-AlexaFluor^488^ stained roots of CCaMKox and *gofCCaMKox* lines colonised by *R. irregularis* at 7 wpi showing smaller abundant septa in intraradical hyphae within *d14l^gofCCaMKox^* lines. At least two independent transformant lines, three plants, three roots, and three arbuscules of each genotype were imaged. One representative image for each genotype of interest is shown, maximum projection of 5-10 Z planes. White channel corresponds to autofluorescence after excitation at 405 nm, green to WGA-AlexaFluor^488^. White arrows highlight hyphal septa. Scale bar, 50 μm.

Notably, at 7 weeks, all *gofCCaMKox* lines had dwarf phenotypes compared to their respective background genotype (Fig. S4A,B, table S5), all *gofCCaMKox* being significantly shorter than WT*^GFP^*, *ccamk*, and *d14l^GFP^* (*p* < 0.01). This phenotype positively correlated with the level of transgene expression (Fig. 1C), in that the highest expressing lines had the strongest growth suppression. In contrast, *CCaMKox* lines did not display any growth defects (Fig. S4A). This phenotype was also independent of mycorrhizal inoculation (Fig. S4B). Together, these results suggest that CCaMK autoactivation rather than overexpression is responsible for the observed effects on growth, and that this is not due to a potential “excessive” engagement with AMF, but other pleiotropic activity of CCaMK autoactivation. However, this dwarf phenotype is relaxed at later stages of vegetative plant growth. When measuring shoot and root biomass at 12 weeks, a time-point where mycorrhizal benefit is evident (increase in shoot and root biomass due to AM inoculation, under low-phosphate fertilisation), only some individual transgenic lines had significantly inferior biomass when compared to WT*^GFP^*: for shoot dry weight, only *ccamk^gofCCaMKox^* #3 and *d14l^gofCCaMKox^* #3 in mock conditions, *d14l^gofCCaMKox^*#2 and 3 under mycorrhizal conditions (Fig. S5, table S6). All WT*^gofCCaMKox^*were statistically equivalent to the WT and WT*^GFP^* control regardless of inoculation. Notably, most *gofCCaMKox* lines showed a mycorrhizal growth benefit, similar to WT and WT*^GFP^* and *CCaMKox* lines, which was absent from *ccamk*, *d14l*, and *d14l^CCaMKox^* lines (Fig. S5, table S6). Thus, ectopic autoactive CCaMK does not appear to limit mycorrhizal benefit or markedly arrest plant growth past early vegetative development.

### Regulation of mesocotyl elongation by D14L does not require CCaMK

Rice D14L is also involved in the control of mesocotyl length (Gutjahr et al., 2015; Kameoka and Kyozuka, 2015), and additional developmental roles of D14L/KAI2 have been described in other species, particularly Arabidopsis, including germination, root hair density, leaf and root architecture (reviewed in Machin et al., 2020; Waters & Nelson, 2023). Given that CCaMK acts downstream of D14L in symbiotic signalling, we investigated whether it similarly functions downstream in a developmental context. To test this, mesocotyl elongation was measured in *CCaMKox* and *gofCCaMKox* lines (Fig. 2, S6A, independent experiment in Fig. S6B, table S7). Neither *CCaMKox* nor *gofCCaMKox* had a significant effect on mesocotyl length in WT, *ccamk*, or *d14l*. It was found that neither *CCaMKox* nor *gofCCaMKox* could rescue mesocotyl development in rice *d14l*, revealing that distinct signalling events occur downstream of D14L for symbiosis and mesocotyl regulation. This is not surprising as nonmycorrhizal plants exhibit D14L-mediated development regulation but have selectively lost a number of key symbiosis genes including *CCaMK* and *CYCLOPS* (Bravo et al., 2016; Radhakrishnan et al., 2020).

### D14L signalling plays a role in arbuscule development and intraradical hyphal growth

Microscopic quantification of AM colonisation in *d14l^gofCCaMKox^* lines led to the observation that arbuscules appeared morphologically smaller compared to WT. The *d14l^gofCCaMKox^* arbuscules appeared smaller and less densely branched compared to WT but still exhibited coarse and fine order branching (Fig. 3A, S7A). The area ratio of arbuscule to cell was determined via confocal laser-scanning microscopy to quantify the arbuscule size phenotype (Fig. 3B, independent experiment in Fig. S7B, table S8). Three independent *d14l^gofCCaMKox^*lines tested resulted in statistically smaller arbuscules compared to all other genotypes (*p* < 0.05, *p* < 0.0001 when compared to WT). WT*^gofCCaMKox^* and *ccamk^gofCCaMKox^* also had a tendency for smaller arbuscules, although this effect was markedly less pronounced than in the *d14l* background, with an average 0.64 ±0.01 for both WT*^gofCCaMKox^* and *ccamk^gofCCaMKox^*, compared to 0.69 ±0.10 in the WT and 0.40 ±0.01 in *d14l^gofCCaMKox^*. This observation that arbuscules were significantly smaller relative to WT in the *d14l* background suggests that D14L signalling has a distinct role in arbuscule development, which has previously not been characterised in rice due to the lack of root colonisation, hence arbuscule formation in the *d14l* mutant (Gutjahr et al., 2015; Choi et al., 2020).

Intraradical hyphal growth in the *d14l^gofCCaMKox^* lines also appeared affected, with abundant septa within the hyphae (Fig. 3C, S8). This was not observed in the WT or *ccamk* background, or for the *CCaMKox* constructs, indicating it is also a consequence of autoactive CCaMK in absence of D14L signalling. AM fungi are typically coenocytic and form septa mainly during senescence or as a damage-repair response, for example to compartmentalise dying hyphae, after germination without a host, or during arbuscule collapse (Smith and Read, 2008; Cargill et al., 2025 and citations therein). The high abundance of septa observed in *d14l^gofCCaMKox^*roots therefore suggests altered intraradical fungal integrity and/or accelerated senescence, possibly reflecting a root environment that is less supportive of fungal growth. This could be associated with the smaller arbuscules, although direction of causality remains unclear: reduced fungal fitness could limit arbuscule maturation, or impaired arbuscule development could constrain carbon delivery and promote septation. Overall, these results suggest that D14L signalling has a role in intraradical fungal growth and arbuscule development that is not bypassed by CSSP activation.

### Gain-of-function CCaMK expands root domains permissive for AM fungal colonisation

Despite total colonisation and arbuscule abundance not being significantly different between all *gofCCaMKox* lines and the WT, the intraradical distribution of AM fungal structures appeared markedly denser in *gofCCaMKox* roots, with more abundant hyphae throughout the cortex, proliferating between most cell layers (figure S1). Consistent with this increased intraradical spread, hyphae in *gofCCaMKox* lines (across all genetic backgrounds) recurrently colonised the meristematic region of the root tip, crossing the quiescent centre (QC) and expanding through the columella, with frequent instances of entry or exit via the root tip (Fig. 4A). We quantified the distance of the closest hyphal tip to the QC in root tips of large lateral roots and found a significant reduction in hyphal distance to QC in *gofCCaMKox* lines, from 0.105 mm ±0.117 mm in the WT to an average of 0.015 mm ±0.04 mm in *gofCCaMKox* lines. In parallel, the frequency of hyphae crossing the QC ranged from 0% in the WT, 8.3% in WT*^GFP^* to 4.3% in *CCaMKox* lines, and increased to 31.9% in *gofCCaMKox* lines on average (Fig. 4B, table S9). Hyphae entering or exiting via the root tip were also more frequent: such events were rare in WT (0% in WT, 12.5% in WT^GFP^) and in *CCaMKox* lines (13.0%), whereas they are surprisingly abundant in *gofCCaMKox* lines (40.3%). Perplexingly, we observed that in *gofCCaMKox* lines AM hyphae could colonise endodermal cells and formed branched structures reminiscent of arbuscules within this layer, a phenotype never observed in WT or *CCaMKox* lines (Fig. 4C, D, S9). Together, these observations indicate that CCaMK autoactivation (and overexpression) expands AM colonisation into domains that are normally non-permissive, thereby proposing CCaMK activation state as a key regulator of root permissiveness across distinct anatomical regions. As these phenotypes were specific to *gofCCaMKox* and not seen in *CCaMKox*, expansion of colonisation domains appears to require CCaMK activation rather than increased CCaMK expression alone.

**Figure 4.**
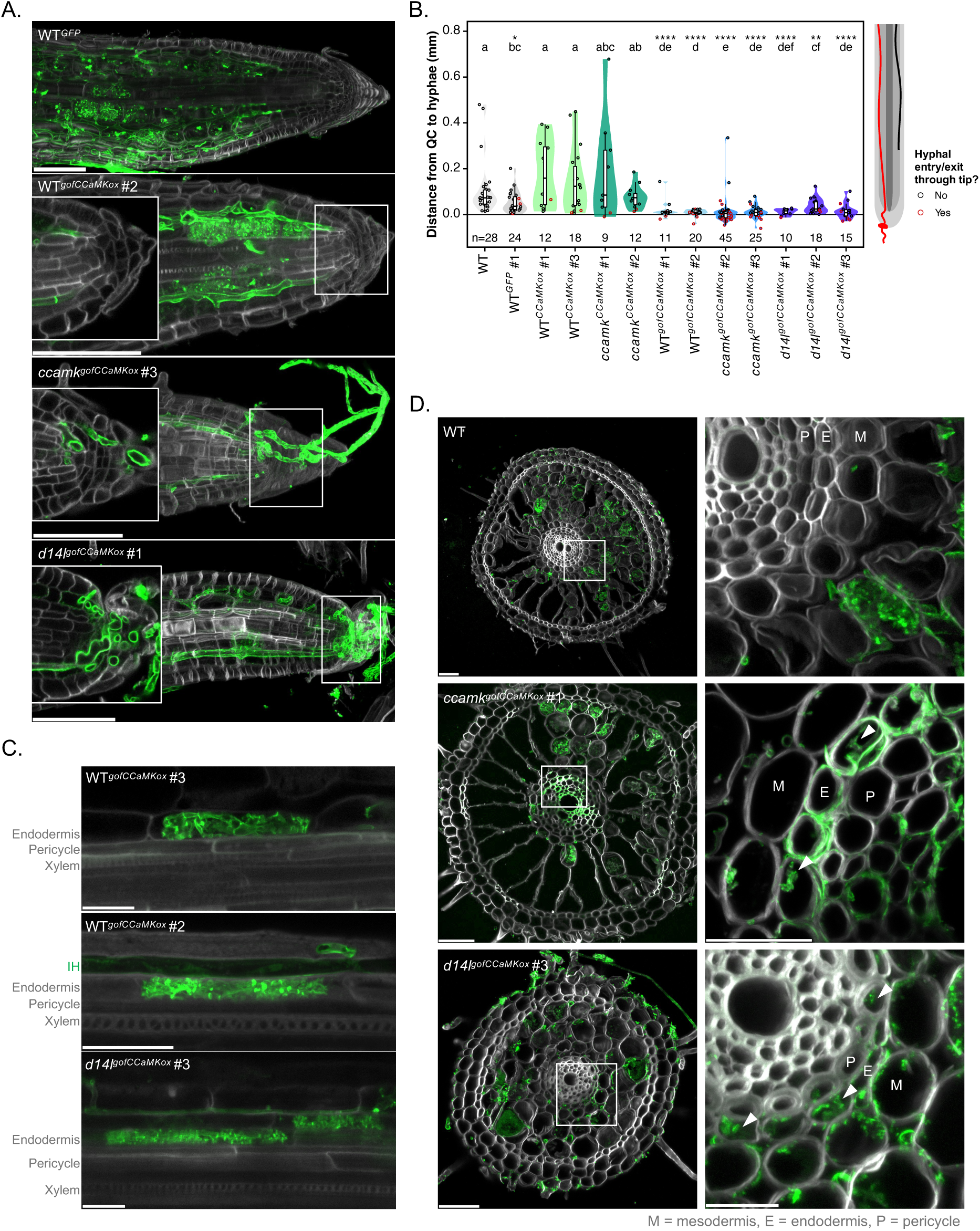
*gofCCaMKox* allows AM colonisation to expand into the root meristem and endodermis. (A) CSLM of WGA-AlexaFluor^488^ stained root tips of *CCaMKox* and gofCCaMKox lines colonised by *R. irregularis* at 7 weeks-post-inoculation (wpi) showing colonisation of meristematic zone in *gofCCaMKox* lines. At least two independent transformant lines, three plants, three roots, and three arbuscules of each genotype were imaged. One representative image for each genotype of interest is shown, maximum projection of 5-10 Z planes. White channel corresponds to autofluorescence after excitation at 405 nm, green to WGA-AlexaFluor^488^. Scale bar, 50 μm. (B) Quantification of distance from hyphal tip to quiescent-centre in roots tips of *CCaMKox* and *gofCCaMKox* lines colonised by *R. irregularis* at 7 wpi, stained either with WGA-AlexaFluor^488^ or Trypan-Blue. Individual datapoints are shown (*n* > 4 root tips, specified on the graph, from ≥ 3 biological replicates). Violin colour represents genotype, point edge colour represents whether hyphae enter or exit root via the tip (red) or not (black). Horizontal line at 0 represents QC, negative numbers indicate hyphae penetrating the lateral root cap or columella. Statistically significant differences are determined by Kruskal-Wallis test, p-value displayed, followed by Pairwise Wilcoxon rank sum test. Letters denote statistically significant differences between groups (*p* < 0.05), asterisks indicate significant differences to WT (*, *p* < 0.05; **, *p* < 0.01; ***, *p* < 0.001; ****, *p* < 0.0001). (C and D) CSLM of WGA-AlexaFluor^488^ stained roots of *gofCCaMKox* lines colonised by *R. irregularis* at 7 wpi showing colonisation of endodermal cells, (C) for longitudinal close-ups, (D) for cross sections. One representative image for each genotype of interest is shown, maximum projection of 3-10 Z planes. White channel corresponds to autofluorescence after excitation at 405 nm, green to WGA-AlexaFluor^488^. Cell layers indicated on the left in (C), within the image in (D), with M for mesodermis (last cortical cell layer before endodermis), E for endodermis, P for pericycle. Colonised endodermal cells highlighted via white arrows in (D). Scale bar, 20 μm.

### Gain-of-function CCaMK leads to transcriptional activation of symbiotic genes even in the absence of AMF

To assess how *CCaMK* expression and activation influence early symbiotic signalling, we profiled transcriptomes of *CCaMKox* and *gofCCaMKox* lines in all three backgrounds at an early stage of colonisation (3 wpi) in low Pi conditions. This time point was chosen to capture early responses before extensive intraradical colonisation becomes established, preventing the masking of decisive early signalling events that underpin symbiosis development.

The colonisation phenotypes at 3 wpi of the lines used for transcriptome analysis were quantified (Fig. S10A). WT*^GFP^* total colonisation was 15.33% ±0.88% compared to 16.67% ±0.88% in WT*^CCaMKox^* and 17.33% ±0.33% in WT*^gofCCaMKox^*. The *ccamk* mutant displayed 1% ±0.57% total colonisation, compared to 5.66% ±2.19% in *ccamk^CCaMKox^* and 16.67% ±0.88% in *ccamk^gofCCaMKox^*. *d14l^GFP^* and *d14l^CCaMKox^* had 0% total colonisation, compared to 16% ±0.57% total colonisation in *d14l^gofCCaMKox^*. To simultaneously determine the effects of *CCaMK* overexpression and gain-of-function in non-mycorrhizal conditions, mock-inoculated plants were grown side-by-side (hence “mock”, versus “myc” for mycorrhizal samples), and lack of fungal contamination confirmed microscopically.

We compared our ‘early interaction’ dataset to published rice root transcriptomes. We used the *smax1* mutant transcriptome (Choi et al., 2020) to identify genes regulated by D14L/SMAX1 signalling, and low/high phosphate mycorrhizal transcriptomes at 6 wpi (∼60% total; ∼50% arbuscules), including of the *phr2* mutant (Das et al., 2022) to compare against highly-colonised root system, and distinguish regulation due to phosphate starvation signalling.

In the WT*^GFP^* control, comparing myc versus mock roots identified 195 up-regulated and 488 down-regulated genes (|FC|>1.5, FDR < 0.05, Fig. 5A). The relatively modest number of induced genes compared to 2160 induced genes at 6 wpi (Das et al., 2022), suggests that sampling captured an early colonisation stage, as expected at 3 wpi (Fig. S10). Consistently, among curated AM-related genes (Das et al., 2022; Table S11), only *AM1*, *AM3*, and *AM11* were significantly induced at this time-point, whereas a larger group of genes was only detected as induced with ∼60% colonisation at 6 wpi (Das et al., 2022), including *PT11*, *RAM1*, *FatM*, *STR1*, *STR2*, *HA1*, *NOPE1*, *PT13*, and *CYCLOPS*; as well as a set of apocarotenoid-related biosynthetic genes and signalling components (*MAX1C*, *CCD7/D17*, *CCD8B/D10*, *DLK2A-C*, *SMAX1*, *NSP2*, *ZAS*) that are known to be regulated by D14L signalling (activated in *smax1*, repressed in *d14l*) (Fig. 5B, S11). Notably, this last gene module appears down-regulated at 3 wpi, including *MAX1C*, *CCD1b*, *CCD7/D17*, *DLK2B* and *DXS2* (Fig. 5B, S11).

**Figure 5.**
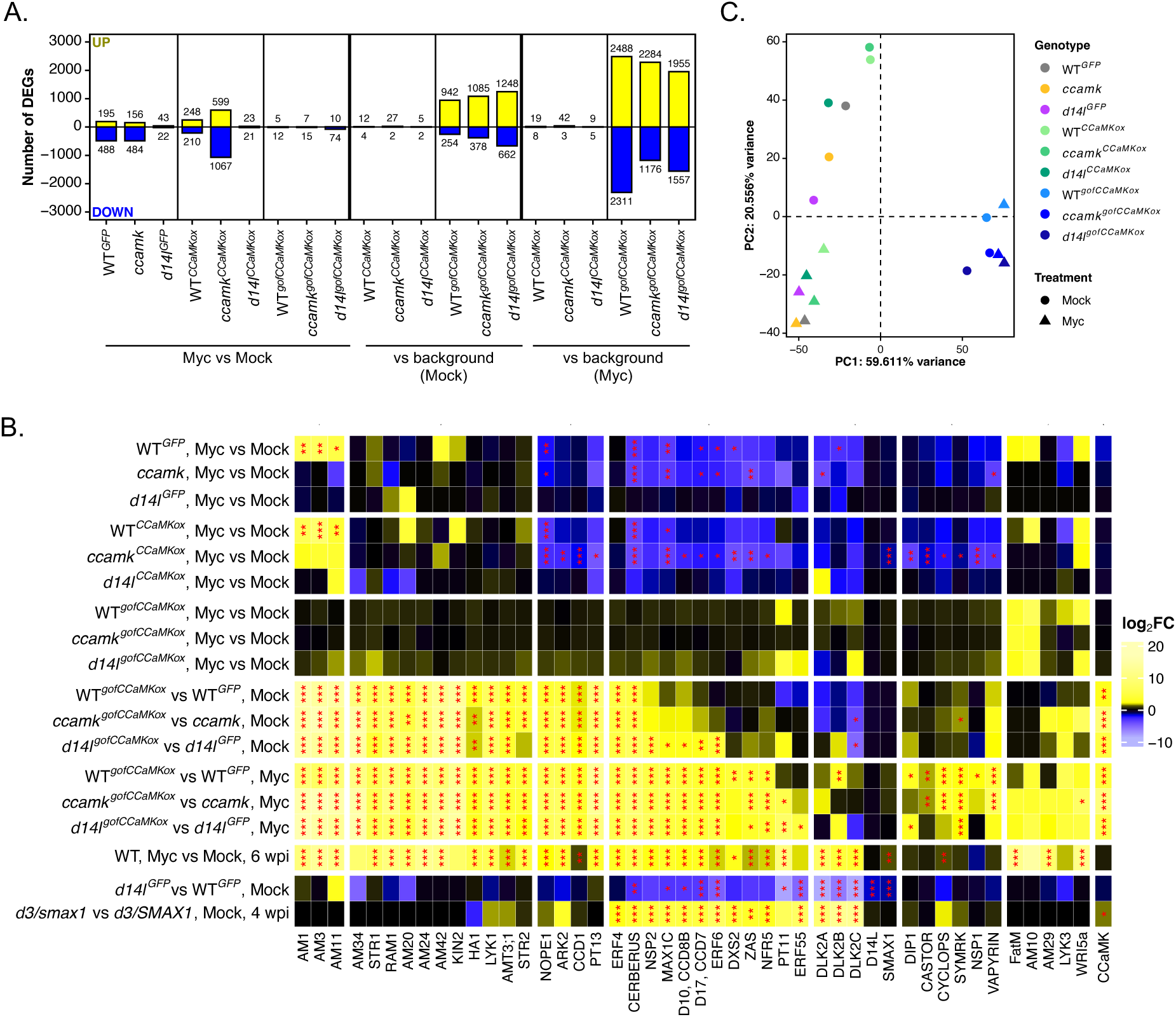
*gofCCaMKox* activates an AM-associated transcriptional programme even in the absence of AM fungi. (A) Number of differentially expressed genes (DEGs), both up-regulated (positive values, yellow) and down-regulated (negative values, blue), for comparisons of interest. Numbers over or below bars indicate the precise number of up or down-regulated genes in each comparison. (B) Principal Components Analysis (PCA) for RNA-seq samples of *CCaMKox* and *gofCCaMKox* lines. Colours indicate genotype, shape indicates treatment (triangles for mycorrhizal samples, squares for non-mycorrhizal samples). Variance-stabilised counts (VST counts) for the 5000 top variable features in the dataset were used to conduct PCA analysis. Only PC1 and PC2 are shown as they together account for more than 50% of the variance. (C) Heatmap of log2FC for comparisons of interest for a selection of genes related to AM symbiosis (subset of the AM gene list from Das et al., 2022; common names shown, gene IDs in table S11). Significance levels as determined by DEG using DESeq2 shown with asterisks (* for *p*-value <0.05, ** for < 0.01, *** for < 0.001). Genes were manually curated into 8 clusters based on expression patterns. Extended version of this heatmap with further AM-related genes in figure S11A.

The *ccamk* mutant responded to inoculation with a DEG magnitude similar to WT (156 up, 484 down), including repression of apocarotenoid-related genes, but lacked induction of early AM markers, consistent with the absence of active CSSP signalling (Fig. 5A, 5B). In contrast, the *d14l* mutant showed a strongly reduced response (43 up, 22 down), lacking both early AM marker induction and the apocarotenoid repression signature, as this module is already repressed in mock conditions in *d14l*, conversely induced for *smax1* (Fig. 5A, 5B, S11), consistent with the lack of pre-symbiotic responsiveness reported previously for *d14l* when exposed to germinating spore exudates (Gutjahr et al., 2015a).

*CCaMKox* lines in WT and *ccamk* backgrounds showed responses to AMF inoculation similar to WT in both DEG magnitude and marker gene activation, whereas *d14l^CCaMKox^* behaved similarly to *d14l*, following phenotypic results (Fig. 5A, 5B, S11A). In contrast, *gofCCaMKox* lines, regardless of genetic background, showed minimal response to AMF inoculation at 3 wpi (17-84 total DEGs). However, when compared to their respective background genotypes (WT*^gofCCaMKox^* to WT*^GFP^*, *d14l^gofCCaMKox^*to *d14l^GFP^* and *ccamk^gofCCaMKox^* to *ccamk*), *gofCCaMKox* lines displayed substantial transcriptome changes already in mock conditions (1196–1910 DEGs) and even more in myc conditions (3460–4799 DEGs), whereas *CCaMKox* lines remained transcriptionally close to their backgrounds (7–45 DEGs; Fig. 5A). PCA supported this interpretation: *gofCCaMKox* expression was the dominant driver of variance, with all *gofCCaMKox* samples clustering together along PC1 (46% of the variance) irrespective of background and inoculation. By contrast, separation of myc versus mock samples for *CCaMKox* lines and background genotypes occurred along PC2 (21% of the variance) (Fig. 5C).

This *gofCCaMKox*-driven transcriptional state included strong induction of many AM symbiosis genes, spanning early markers (*AM1*, *AM3*, *AM11*) and genes typically induced later (e.g., *STR1*, *RAM1*, *LYK1*, *PT13*, *AM42*, *AM34*, *ARK2*) (Fig. 5B, S11). Notably, symbiotic gene activation in the *gofCCaMKox* lines was almost identical between myc and mock samples, indicating that *gofCCaMKox* can activate symbiotic gene expression even in the absence of AMF, as reported in *Lotus japonicus* (Takeda et al., 2015). However, some genes (including *VAPYRIN*, *SYMRK*, and *CYCLOPS*) and several D14L-regulated apocarotenoid-related genes (*NSP2*, *MAX1C*, *D10/CCD8B*, *D17/CCD7*, *DXS2*, *ZAS*) were induced only in *gofCCaMKox* myc samples, suggesting they require additional inputs associated with intraradical colonisation and are not directly activated by autoactive CCaMK alone. Overall, the muted myc-versus-mock response in *gofCCaMKox* lines at 3 wpi reflects a basal activation of AM-associated gene expression already in mock conditions, driven by CCaMK autoactivation rather than just CCaMK overexpression.

### Overexpression of gain-of-function CCaMK drives excessive regulation of symbiotic and non-symbiotic processes, potentially via AP2 transcription factors

The magnitude of transcriptome alteration in *gofCCaMKox* lines, vastly surpassing the response to mycorrhizal inoculation at this time-point, and even that of a highly colonised root system at 6 wpi (2160 up, 340 down; Das et al., 2022; Fig. S12A), led us to further investigate this transcriptional signature. Particularly notable was the high volume of down-regulated genes in WT*^gofCCaMKox^* versus WT*^GFP^* for mycorrhizal samples of 2311 repressed genes, of which only 93 were shared with the AM-repressed transcriptome at 6 wpi (Fig. S12A). We hypothesised CCaMK activity in non-native tissues or cell types, alongside lack of autoinhibitory control, might drive gene modules regulated by mycorrhizal colonisation to excessive activation or repression, and even regulate other biological processes normally not affected by mycorrhizal colonisation. This would be consistent with the pleiotropic growth penalties observed in *gofCCaMKox* lines (Fig. S4) and could reflect the reported expansion of AMF colonisation domains to normally non-permissive tissues or cell-types (meristem, endodermal cells).

To define a common *gofCCaMKox* signature, we intersected DEG sets of each *gofCCaMKox* line against its background genotype, separately for mock and myc conditions. This yielded 564 induced / 51 repressed shared DEGs in mock and 1228 induced / 490 repressed shared DEGs in myc (Fig. 6A; Table S12). GO enrichment analyses (Fig. 6B, C) highlighted expected symbiosis-related terms among induced genes, as well as enrichment for oxalate metabolism in both conditions; additional enrichments in myc included gibberellin, acetyl-CoA, and carotenoid biosynthetic processes. Repressed genes were enriched for iron starvation in mock, while in myc they additionally included response to anoxia, metal ion transport, amino acid transport, and auxin-activated signalling (Fig. 6B, C). Notably, some induced processes (e.g. gibberellin biosynthesis and carbon metabolism) are also observed during WT mycorrhizal colonisation at 6 wpi, whereas repression of modules such as auxin signalling and amino acid/metal transport is less characteristic of later WT symbiosis. This supports the idea that CCaMK autoactivation results in disproportionate physiological reprogramming, which could contribute to growth penalties.

**Figure 6.**
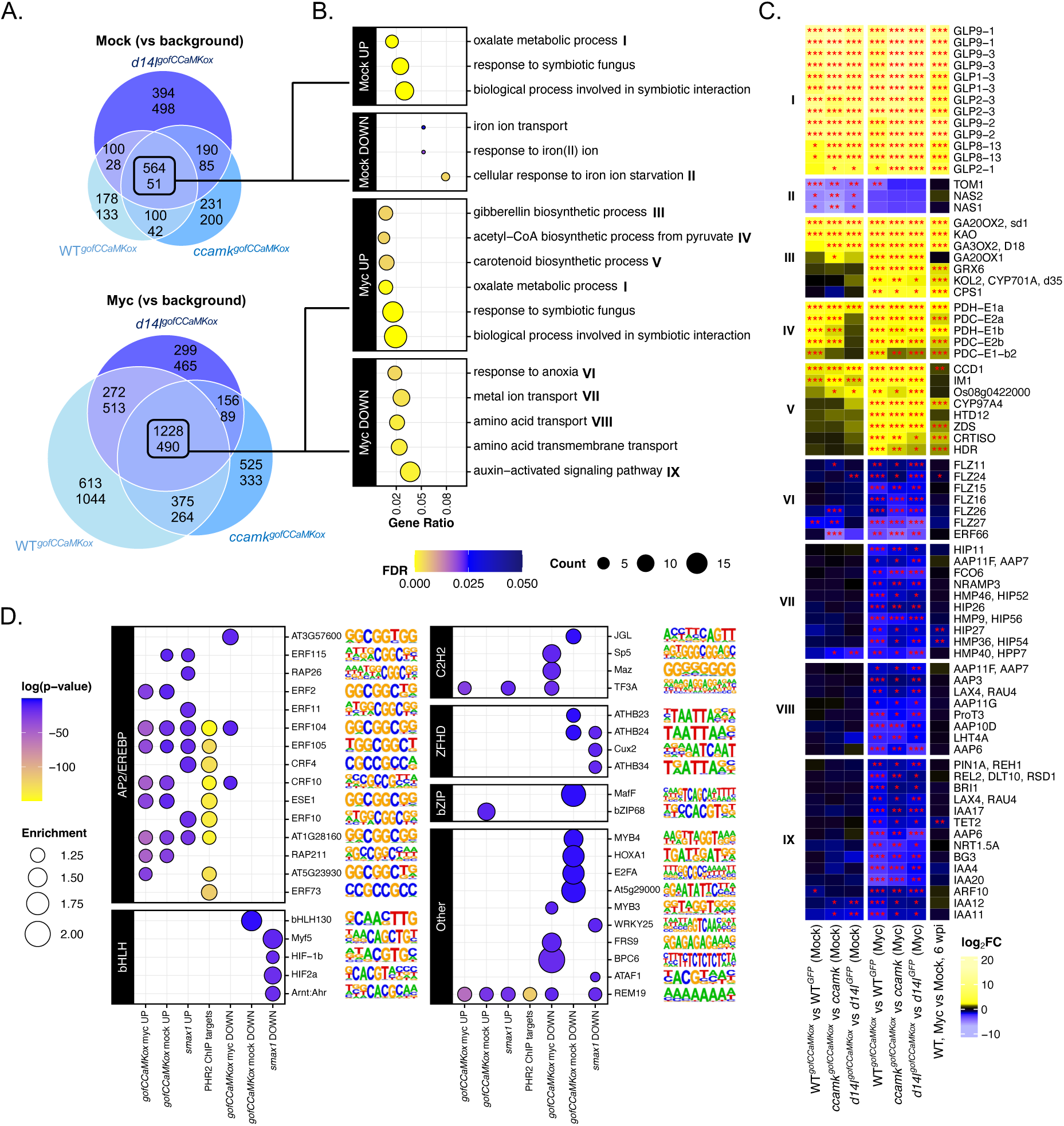
CCaMK autoactivation drives broad ectopic reprogramming with enrichment for AP2/EREBP regulatory signatures. (A) Venn diagram of overlaps between DEGs for *gofCCaMKox* lines in different backgrounds, in non-mycorrhizal and mycorrhizal samples. Upper number for up-regulated genes, lower number for down-regulated genes. (B) GO term enrichment for common DEGs on all *gofCCaMKox* lines in mock and myc conditions, top three to six (based on enrichment values) enriched GO terms by gene ratio for each comparison shown. Colour scale indicates significance as q-value or FDR; dot size proportional to the number of genes in each set included in the GO term; x-axis indicates the ratio of genes up-regulated in each set included in the GO term to the total number of genes annotated with the GO term. (C) Heatmap of log2FC for comparisons of interest (*gofCCaMKox* lines against their background in mock and myc conditions, myc versus mock at 6 weeks-post-inoculation from Das et al., 2022) for genes differentially regulated commonly by *gofCCaMKox* and included in a GO term of interest from panel B. Significance levels as determined by DEG using DESeq2 shown with asterisks (* for *p*-value <0.05, ** for < 0.01, *** for < 0.001). (D) Motif enrichment analysis results for promoters (defined as 1kb upstream of ATG) of selected gene sets including common DEGs for *gofCCaMKox* lines, *smax1* up/down genes from Choi et al., 2020; and PHR2 ChIP targets from Das et al., 2022. Colour scale indicates significance as log(*p*-value); dot size proportional to enrichment value of each motif in each gene set. Separate boxes for motifs corresponding to distinct transcription factor families, motifs and their associated transcription factor indicated on the right.

We next performed motif enrichment analysis on promoters of the shared gene sets to identify candidate transcription factors acting downstream of CCaMK. Shared induced genes (in both mock and myc) were strongly enriched for AP2/EREBP transcription factor binding sites (Fig. 6D; Table S13). This is consistent with current models in which CCaMK–CYCLOPS activates RAM1, which acts alongside and/or through AP2 regulators such as WRI5 and CBX1 to induce symbiotic genes (Luginbuehl et al., 2017; Jiang et al., 2018; Xue et al., 2018; Leng et al., 2023; Zhang et al., 2023). The *CYC-RE* binding motif of CYCLOPS was not detected as a known or de novo enriched motif (Table S13). The enrichment for AP2/EREBP motifs was also observed among *smax1* up-regulated genes (Choi et al., 2020) and PHR2 ChIP-seq targets (Das et al., 2022) (Fig. 6D), consistent with overlap among nutrient- and symbiosis-regulated gene modules (Fig. S12B). In contrast, down-regulated genes showed weaker and more condition-specific motif enrichments (Fig. 6D), suggesting they may be more indirectly regulated. Overall, these analyses are consistent with the hypothesis that CCaMK-driven transcriptional activation is channelled through AP2/EREBP-centred regulatory logic during pre-symbiosis and intraradical colonisation. Notably, three AP2-domain containing Ethylene Response Factors; *ERF4* (*Os12g0582900*), *ERF6* (*Os07g0575000*) and *ERF55* (*Os06g0160500*); were part of the *smax1* up-regulated gene list (Choi et al., 2020), of which *ERF4* and *ERF6* are also induced by *gofCCaMKox* (Fig. 5C), marking them as potential candidates to mediate this transcriptional activation.

### D14L regulates nutrient- and immunity-related gene modules independently of CCaMK

We then interrogated the dataset for transcriptional signatures that could be associated with the change in fungal morphology observed in *d14l^gofCCaMKox^*. Comparing *d14l^gofCCaMKox^* to WT*^gofCCaMKox^* identified 31 induced / 44 repressed genes in myc samples and 144 induced / 42 repressed genes in mock, with 23 induced / 17 repressed genes shared irrespective of fungal colonisation (Fig. 7A; Table S14). As expected, D14L signalling markers *DLK2B*, *DLK2C*, *SMAX1*, and *D14L* were repressed in both conditions, consistent with the RT-qPCR data and supporting that these genes are controlled by D14L/SMAX1 signalling independent of autoactive CCaMK. In contrast, several genes typically up-regulated in *smax1* and repressed in *d14l* (including strigolactone biosynthetic genes such as *CCD7*, *CCD8*, *D27*, as well as *NSP2*) were no longer repressed in *d14l^gofCCaMKox^*, suggesting that autoactive CCaMK can override D14L-dependent repression for this gene module (Fig. 5B, 7B).

**Figure 7.**
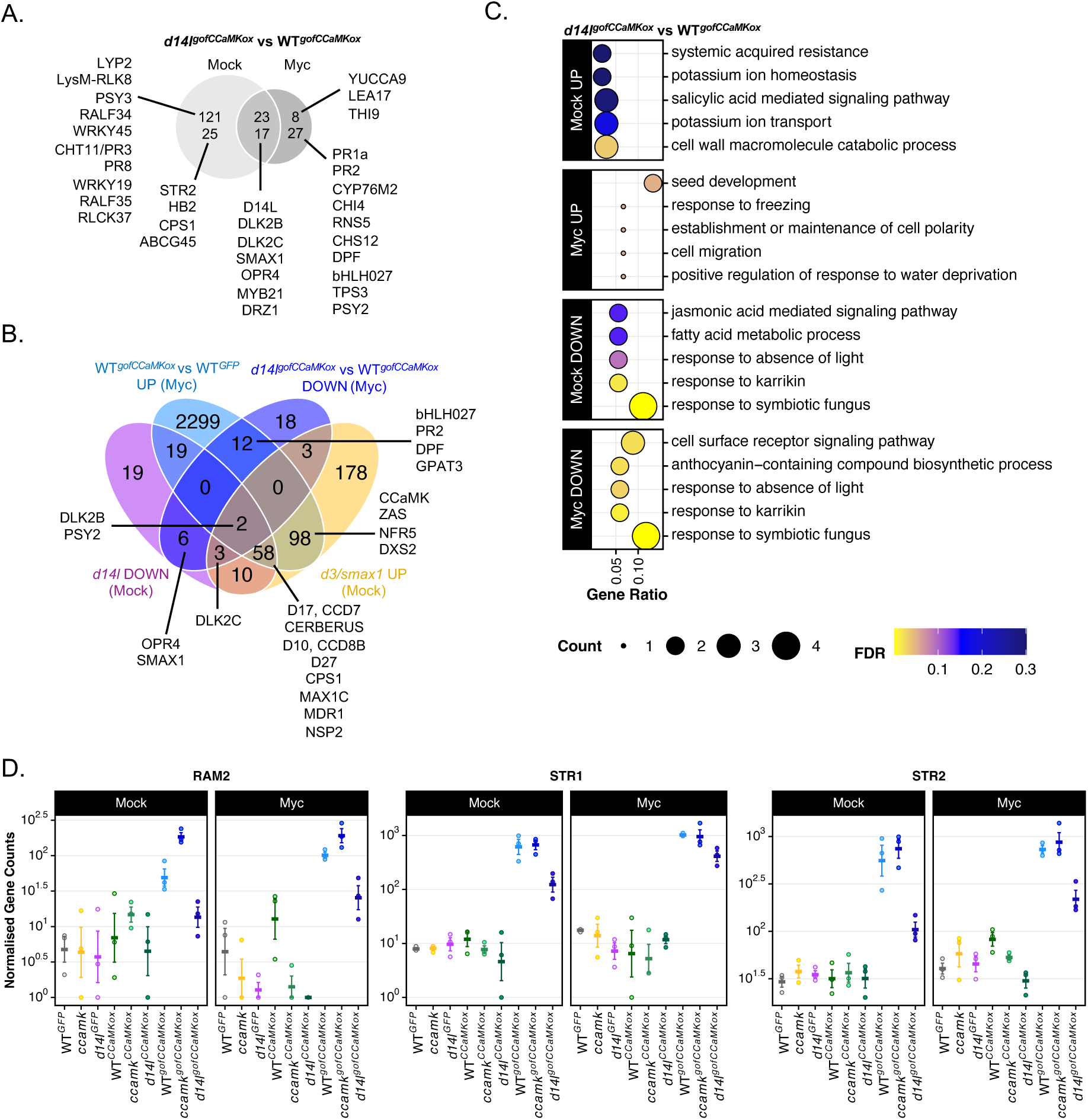
D14L regulates a CCaMK-independent subset of nutrient- and immunity-related genes during symbiosis. (A) Venn diagram of overlaps between DEGs for *d14l^gofCCaMKox^*vs WT*^gofCCaMKox^* in non-mycorrhizal and mycorrhizal samples. Upper number for up-regulated genes, lower number for down-regulated genes. Genes of interest highlighted. (B) GO term enrichment for *d14l^gofCCaMKox^*vs WT*^gofCCaMKox^* DEGs in mock and myc conditions, top five enriched GO terms by gene ratio for each comparison shown. Colour scale indicates significance as q-value or FDR; dot size proportional to the number of genes in each set included in the GO term; x-axis indicates the ratio of genes up-regulated in each set included in the GO term to the total number of genes annotated with the GO term. (C) Venn diagram of overlaps between down-regulated genes for *d14l^gofCCaMKox^*vs WT*^gofCCaMKox^* in myc samples, down-regulated genes for *d14l^GFP^* vs WT*^GFP^* in mock samples, up-regulated genes for WT*^gofCCaMKox^* vs WT*^GFP^* in myc samples, and *d3/smax1* vs *smax1* in mock samples from Choi et al., 2020. Genes of interest highlighted. (D) Normalised gene counts for *RAM2*, *STR1* and *STR2*.

Beyond pathway markers, down-regulated genes in *d14l^gofCCaMKox^* were enriched for immunity and stress-related functions, supported by GO term enrichment, and included several transcription factors (Fig. 7A, C, table S14). A MYB TF (*JAmyb/MYB2*1), the putative oxo-phytodienoic acid reductase *OPR4*, and the zinc finger protein *DRZ1/ZFP36*, appear commonly downregulated in mock and myc conditions. In myc samples, down-regulated genes also included defence-related *PR1a* and *PR2/GNS4*, chalcone synthase *CHS12*, phytoene synthase *PSY2*, terpene synthase *TPS3,* chitinase *CHI4*, two LEA proteins (*LEA2*, *LEA33*), and a bHLH TF involved in regulation of diterpenoid phytoalexin biosynthesis, *DPF*. In mock conditions, the kaurene synthase *CPS1* and notably *STR2* were repressed (Fig. 7A; Table S14). The other symbiotic half-sized ABCG transporter, *STR1*, and the arbuscule-specific glycerol-3-phosphate acyltransferase *RAM2* showed similar (though not significant) trends. Additional nutrient signalling components or transporters/regulators (*PK*, *NFP4.5*, *VIP1*, *SPX5*, *PT9*, *PT10*, *SPX1*, *PT6*) followed a similar pattern (Fig. 7D, S12C; Table S15). Notably, mutants in nutrient transporters and signalling components, such as rice *str1*, *str2* and *ram2* mutants, exhibit stunted arbuscules (Gutjahr et al., 2012; Liu et al., 2022). Thus, these expression changes may contribute to, or reflect, the reduced arbuscule size and altered fungal fitness in *d14l^gofCCaMKox^*. Likewise, reduced induction of immunity/stress signatures could influence fungal accommodation but could also be a consequence of altered arbuscule maturation dynamics, as these signatures can be enriched in the translatome at particular stages of arbuscule development and collapse (Chancellor and Ferreras-Garrucho et al., 2026). Overall, despite the relatively small number of affected genes, these results support that, in the context of active CSSP signalling driven by autoactive CCaMK, loss of D14L signalling is associated with specific alterations in nutrient- and immunity-related gene modules that could underlie, or respond to, the impaired intraradical growth and arbuscule development phenotypes.

## Discussion

Here, we assessed the epistatic relationship between D14L signalling and the common symbiosis signalling pathway (CSSP). Prior evidence that CSSP components are transcriptionally upregulated in *smax1* (Choi et al., 2020), together with reduced CO/LCO-induced Ca^2+^-spiking in rice *d14l* (Li et al., 2022), placed D14L upstream of the CSSP. Consistent with this, autoactive CCaMK (*gofCCaMKox*) restored AM symbiosis in *d14l*, whereas constitutive CCaMK overexpression failed to do so. These data support a model in which a key output of D14L signalling, rather than only inducing expression of CCaMK, is to enable CCaMK activation and downstream transcriptional reprogramming required for AM colonisation.

RNA sequencing of *CCaMKox* and *gofCCaMKox* lines provides molecular support for this model, resolving how CCaMK activation shapes the symbiotic transcriptome. Introduction of *gofCCaMKox* induced expression of many AM-associated genes (e.g., *AM1*, *AM3*, *RAM1*, *STR1*, *STR2*, *HA1*, *AMT3;1*) in non-mycorrhizal roots across genetic backgrounds, reaching levels comparable to those observed later during colonisation (Das et al., 2022). However, a subset of AM genes, featuring CSSP components *CASTOR*, *POLLUX*, *CYCLOPS* and *SYMRK,* were only activated in *gofCCaMKox* lines in AMF-inoculated roots, indicating that additional mycorrhiza-specific inputs, potentially linked to fungal signal perception or intraradical accommodation, are required for their induction. This may reflect a regulatory mechanism in which the CSSP cannot fully self-activate in the absence of fungal cues, potentially as a safeguard against runaway positive feedback during AM signalling. Within these, a gene module including strigolactone biosynthetic genes *CCD7* and *CCD8*, as well as *NSP2*, *NFR5*, *PT11*, and *ZAS*, were differentially expressed in the *d14l* and *smax1* mutants. Conversely, *SMAX1*, *D14L*, and *DLK2s* were exclusively regulated by D14L/SMAX1, not affected by *gofCCaMKox*. Together, these patterns separate AM-related genes into (i) responsive to CCaMK activation, (ii) requiring CCaMK plus additional mycorrhizal cues, (iii) influenced by CCaMK, mycorrhizal cues and D14L/SMAX1 signalling, and (iv) predominantly controlled by D14L/SMAX1 independently of CCaMK.

Following the existence of CCaMK-independent transcriptional targets of D14L signalling, a key finding of this study is that the symbiotic role of D14L extends beyond enabling CSSP activation, as arbuscules in *d14l^gofCCaMKox^*were smaller, and intraradical hyphae harboured abundant septa. This phenotype, inaccessible in rice *d14l* due to the near-complete colonisation block (Gutjahr et al., 2015), points to a previously underappreciated contribution of D14L signalling to intraradical development and/or arbuscule maturation. Consistent with impaired arbuscule function, arbuscule-specific *PT11* and *ARK2* were shown to be up-regulated in the *smax1* mutant (Choi et al., 2020), and the transcriptome of *d14l^gofCCaMKox^*showed reduced induction of multiple nutrient transport and arbuscule-associated genes (including *STR1/STR2* and others), as well as changes in immunity-related genes and transcription factors. Whether these transcriptional differences are causal drivers of defective arbuscule development or reflect downstream consequences of altered fungal growth remains unresolved, but they collectively argue that D14L has CCaMK-independent outputs that become critical during the intraradical accommodation of AMF.

While these data place CCaMK activation downstream of D14L for AM establishment, the mechanism linking D14L/SMAX1 to CCaMK activation remains unclear. A parsimonious model is that D14L signalling promotes the expression and/or competence of receptors and signalling components required for robust symbiotic Ca^2+^-spiking, thereby pushing CCaMK activation above a critical threshold. More generally, the inability of *gofCCaMKox* to rescue the *d14l* mesocotyl phenotype, together with identification of genes regulated by D14L/SMAX1 but not by CCaMK, supports bifurcation of D14L signalling into developmental and symbiotic branches. Following insights from the paralogous D14 signalling pathway (Jiang et al., 2013; Wang et al., 2015; Ma et al., 2017; Tang and Chu, 2020; Wang et al., 2020), identifying the transcriptional regulators and co-repressor modules operating downstream of SMAX1 (Soundappan et al., 2015; Zheng et al., 2020) in each context remains an important direction, particularly given the large number of transcription factors misregulated in *smax1* (Choi et al., 2020). Notably, these include several AP2-domain containing ERFs, some of which are also regulated by *gofCCaMKox* (*ERF4*, *ERF6*). Consistent with this, genes induced by *gofCCaMKox* were enriched for AP2 transcription factor binding motifs. Similar AP2 motif enrichment in the upregulated transcriptome of the *smax1* mutant (Choi et al., 2020) further supports the epistasis CCaMK and D14L/SMAX1. Overall, while motif enrichment does not establish direct regulation, CCaMK activation engages a transcriptional architecture with a strong AP2 participation, integrating symbiotic and nutrient-responsive gene expression.

Apart from symbiotic gene regulation, colonisation phenotypes of *gofCCaMKox* lines point to a role for CCaMK activation in defining which root domains are permissive to AM fungi. In wild-type angiosperm roots, the meristematic zone and endodermis are typically restrictive to intraradical AM colonisation. Here, *gofCCaMKox* enabled extensive colonisation in the root tip meristem and arbuscule formation in endodermal cells. This expansion is consistent with CCaMK as a central decoder of symbiotic Ca^2+^ signalling even in later stages of colonisation (Sieberer et al., 2012), and suggests that the spatial pattern of CCaMK expression/activation defines anatomical boundaries for colonisation.

In summary, this work supports a model in which D14L signalling promotes AM symbiosis in rice by enabling CCaMK activation, a decisive step for launching the symbiotic transcriptional programme to accommodate the fungus in the root tissue. At the same time, our findings indicate that D14L is required for adequate fungal development and arbuscule maturation through outputs that are not bypassed by constitutive CCaMK activity. More broadly, CCaMK activation emerges not only as a molecular “on switch” for symbiotic gene expression, but also as a key determinant of tissue permissiveness, defining within which tissue domain AM fungi can proliferate and form arbuscules. Together, these conclusions refine our knowledge of AM signalling by distinguishing the converging and diverging roles of two of the central pathways for its regulation, D14L and CSSP, at the genetic, transcriptomic, and anatomical levels.

## Materials and methods

### Plant genetic material

All experiments were conducted using *Oryza sativa* subsp. *japonica* cv. Nipponbare as a background genotype (hereafter WT, unless otherwise indicated). The rice *d14l-3* mutant (hereafter *d14l*) was previously produced in the Nipponbare background using CRISPR/Cas9 technology that introduced a T insertion in the second exon, generating an early stop codon at amino acid 175, resulting in the loss of 96 amino acids from D14L (Choi et al., 2020). Homozygous seed from *ccamk-2* (NF8513) (hereafter *ccamk*) Tos17 (Transposon of *Oryza sativa* 17) insertional mutant line carrying a Tos17 insertion (4,114 bp) in the third intron were previously obtained from the National Institute of Agrobiological Sciences (NIAS, Japan) (Miyao et al., 2003)(Chen et al., 2007; Gutjahr et al., 2008). The transgenic lines in these three backgrounds (*WT*, *ccamk*, *d14l*) employed in this study, detailed in table S1 were generated using Golden Gate Cloning and rice calli transformation.

### Golden Gate cloning

Constructs were generated using Golden Gate cloning (Engler et al., 2009; Weber et al., 2011). Level 0 modules were designed using Benchling (Biology Software) (https://benchling.com). Firstly, the nucleotide sequences of separate modules were domesticated to remove BsaI and BpiI restriction sites considering rice codon usage. Sequences were flanked by 5’ and 3’ end linkers that contain restriction sites for Golden Gate cloning. Golden Gate cloning was simulated *in silico* using Benchling prior to module synthesis to confirm correct design. Gene module synthesis was carried out using GeneArt^TM^ Gene Synthesis (Invitrogen, Carlsbad, USA) or generated from existing level 0 vectors by PCR and transformed into level 0 vectors containing spectinomycin (Spec) antibiotic selection cassettes. Level 0 modules were assembled into the EC47732_pL1V-F1 vector backbone. Level 1 modules were assembled into the EC60606_pL2V-GW level 2 vector backbone flanked by Gateway attL1 and L2 sites and the end linker Golden Gate vector EC41722_ELE1. The final level 2 constructs were recombined into the binary destination vector pEW343-R1R2 containing a Hyg selection cassette for *Agrobacterium tumefaciens*-mediated rice callus transformation.

### Rice calli transformation

Resulting T-DNA plasmids were introduced into *Oryza sativa* subsp. *japonica* cv. Nipponbare embryogenic calli, derived from mature seed embryos, via *Agrobacterium tumefaciens-*mediated transformation. Embryogenic callus was generated by plating surface-sterilised mature seed, with embryo axes removed, on N6DT medium (3.95 g/L N6 basal salts, 30g/L sucrose, 300mg/L casein hydrolysate, 100mg/L myo-inositol, 2878mg/L proline, 0.5mg/L nicotinic acid, 0.5mg/L pyridoxine HCl, 1mg/L thiamine HCl, 37.3 mg/L Na2EDTA, 27.8 mg/L FeSO4, 2mg/L 2,4-D Na salt, 150mg/L Timentin, 4g/L Gelrite, pH5.8). Plates were sealed with Parafilm and cultured in the dark at 28°C for 21 days, after which time callus was cut into 2-4mm pieces, plated on fresh N6DT, and cultured as before for a further 4 days. Preparation of *Agrobacterium* strain EHA105 containing the binary constructs and transformation of the rice callus pieces was carried out as previously described (Choi et al., 2020). Successful transformation was verified by PCR genotyping. In the subsequent T1 generation, plants carrying the transgene and Hyg were chosen for seed propagation. The T2 generation was used for experimental work including AM colonisation, mesocotyl elongation, microscopy, RT-qPCR and RNA-sequencing. Primers for genotyping are listed in table S2.

### Plant growth conditions and AM fungi inoculation

Seeds of *Oryza sativa* subsp. *japonica* cv. Nipponbare were surface-sterilised briefly in 70% (*v/v*) ethanol, then for 20 min in 3% (*v/v*) sodium hypochlorite. Imbibed seeds were germinated on 0.9% (*w/v*) bactoagar at 30°C for 7 days. Pre-germinated seedlings were transferred into grey or black opaque cones (12 cm depth, 2.5 cm diameter) for all assays (AM colonisation quantification, RT-qPCR, RNA-seq) except for biomass measurements, for which they were transferred into containing sterile quartz sand in walk-in growth chambers with photoperiod of 12-hour day-night cycle at 28/20°C, 65% relative humidity under LED illumination at 300 μmol/μm^2^s. The AM fungal model species *Rhizophagus irregularis* (DAOM197198) was employed for all inoculation assays. The source of AM fungi inoculum was a mix of *Agrobacterium rhizogenes* transformed carrot hairy root cultures (*Daucus carota* L.) (Bécard and Fortin, 1988) from which spores were extracted as previously described (Gutjahr et al., 2008). Plants were inoculated with 300 spores of *R. irregularis* per plant, as described previously (Gutjahr et al., 2008; Yang et al., 2012; Chiu et al., 2018). diH2O without spores was used for mock treatments. Plants were watered three times weekly for the first two weeks post-inoculation (wpi), thereafter fertilised twice a week with half Hoagland solution (25 µM Pi) and 0.01% (*w/v*) Sequestrene Rapid (Syngenta). These growth conditions were demonstrated previously to promote efficient and equal mycorrhization (Gutjahr et al., 2008; Chiu et al., 2022).

### Root staining and mycorrhizal quantification

Roots were harvested at 7 wpi. Trypan blue (Sigma-Aldrich) staining and mycorrhizal colonisation of the different genotypes were quantified as described previously (Gutjahr et al., 2008; Chiu et al., 2022). Fungal colonisation was quantified by counting extraradical hyphae, intraradical hyphae, hyphopodia, arbuscules, vesicles, and spores along 100 different visual fields under a 20× magnification objective using DM750 Microscope (Leica, Germany) with GXCAM HICHROME-HR4 LITE C-Mount Camera (GT-Vision, UK) attached to a computer with GX Capture software (GT-Vision, UK). Colonisation as percentage of total root length for each structure was calculated. Representative images were captured. Image analysis was carried our using Fiji under ImageJ software license (Schindelin et al., 2012). For hyphal tip to QC measurements, captured images of colonised root tips were imported into ImageJ, and the segmented line tool was used to measure the distance from the closest hyphal tip to the predicted quiescent centre location.

### WGA-Alexafluor staining for fungal morphology analysis and arbuscule size quantification

Roots were harvested as above into 50% EtOH and incubated at 4°C overnight. Then they were incubated in 20% (w/v) KOH at RT for 2-3 days, rinsed three times with diH_2_O and incubated for 2h in 0.1 M HCl. Roots were rinsed twice with diH_2_O and once with phosphate-buffered saline (PBS; 137 mM NaCl, 2.7 mM KCl, 10 mM Na_2_HPO_4_, 1.8 mM KH_2_PO_4_, pH 7.4) solution before being incubated with 0.2 µg/ml WGA-Alexa Fluor™ 488 (Invitrogen, USA) solution in PBS at 4°C in the dark for at least two weeks. For cross sections, live roots were harvested, cleaned with diH_2_O, then hand sectioned using a single-edged razor blade (Wilkinson, UK) in a 0.4 μg/mL WGA-Alexa Fluor™ 488 (Invitrogen, USA) solution in PBS, then incubated for 5 min at RT. Stained roots were then imaged using CSLM on a Leica Stellaris 8 FALCON FLIM (Leica, Germany). WGA-Alexa Fluor™ 488 was detected using the white light laser (WLL) with an excitation wavelength of 488 nm, and emitted wavelengths collected at 492-533 nm. Plant cell-wall autofluorescence to visualise cell boundaries was detected using the UV 405 nm laser and emitted wavelengths collected at 410-445 nm. Roots were imaged using the 40× water immersion objective, or the 20x dry objective for cross sections. Single plane images at the point of maximal arbuscule expansion or Z-stacks were acquired with a line average of 4 and dimension of 1024 × 1024 or 512 pixels and speed of 400 Hz. Image analysis was carried out using Fiji under ImageJ software license (Schindelin et al., 2012). For the arbuscule/cell ratio measurements, the freehand selection tool was used to individualise cells and arbuscules and the area of selection was measured. A minimum of 50 arbuscules from three independent plants for each genotype studied was measured (Carotenuto & Genre, 2020; Montero & Paszkowski, 2022). For hyphal tip to QC measurements, the segmented line tool was used to measure the distance from the closest hyphal tip to the predicted quiescent centre location in maximum projection of Z-stacks.

### Mesocotyl elongation quantification

*O. sativa* seeds were dehusked, surface-sterilised as described previously, and sowed in two rows in square 12 cm petri-dishes placed vertically upright containing 0.6% (w/v) Bacto^TM^ Agar (sealed with Micropore^TM^ tape), then placed in a 30°C incubator in the dark. Mesocotyls of 7-day-old seedlings were imaged using Leica S9D Stereomicroscope (Leica, Germany) connected to a GXCAM HiChrome HR4K Lite Camera (GT Vision, UK). Mesocotyl length, distance between the points of seminal root emergence and the coleoptilar node, was measured using Fiji under ImageJ software license (Schindelin et al., 2012) using the freehand line selection tool.

### RNA extraction, cDNA synthesis, and gene expression analysis

Roots harvested were frozen in liquid nitrogen and stored at −80°C until used for RNA extraction. Root tissues were homogenised with metal beads using Geno/Grinder® (SPEX SamplePrep, USA) at 1500 strokes/minute for 1 min until fine powder. RNA was extracted from ground tissue with TRIzol reagent (Invitrogen, USA) with chloroform washes and isopropanol precipitation, and residual DNA removed by treatment with DNaseI (Invitrogen, USA) following manufacturer’s instructions. RNA integrity and purity were assessed by presence of clear 28S and 18S rRNA bands following gel electrophoresis, and quantity determined by NanoDrop 2000 Spectrophotometer (Thermo-Fisher Scientific, USA). cDNA synthesis was conducted as described previously (Chiu et al., 2022) where reverse transcription was performed on 1.1 μg of RNA using Superscript IV reverse transcriptase and oligo(dT)15 primers following manufacturer’s instructions. Absence of contaminating genomic DNA was confirmed by performing PCR with primers on two exons flanking a spliced intron in GAPDH to yield a smaller product following electrophoresis on a 0.8% (w/v) agarose gel, with gDNA sample as positive control. Quantitative polymerase chain reaction (qPCR) was performed as described previously (Gutjahr et al., 2008; Chiu et al., 2022), using SYBR Green Fluorophore on C1000 Thermal Cycler with CFX384 real-time detection system (Bio-Rad, USA). Specific primers (table S2) were used for *CYCLOPHILIN2*, *ACTIN*, and *UBIQUITIN* as housekeeping genes, and for the AM and D14L/SMAX1 signalling marker genes of interest (figure S9). Gene expression values were normalised to the geometric mean of the three reference genes and transformed using 2^−(*Ct-Geomean Ct*)^, where Geomean Ct is the geometric mean of the reference genes’ Ct values; expression values are the mean of three technical replicates and outliers were excluded prior to analysis.

### RNA-Seq library preparation, sequencing, and data analyses

RNA extraction and quality controls were conducted as previously described. RNA samples were diluted to an equal concentration and sent for Illumina NovaSeq 6000 sequencing to Novogene (Cambridge, UK) with a sequencing depth of 3 Gb (10 million reads). Raw reads were pseudo-aligned to the *O. sativa* ssp. *japonica* cv. Nipponbare reference transcriptome (Os-Nipponbare-Reference-IRGSP-1.0.52)(Kawahara et al., 2013) using Kallisto v0.46.1 (Bray et al., 2016) with default parameters, to obtain count estimates. An average 89.61% of the reads were aligned to the rice transcriptome. The count estimates were normalised, filtered (keeping only genes with more than 10 counts among all samples), and subjected to pairwise differential expression analysis between conditions using the R package DESeq2 v1.40.2 (Love et al., 2014), with a threshold of fold-change ≥ 1.5 or ≤ −1.5 and Benjamini-Hochberg false discovery rate corrected *P*-value < 0.05. Gene Ontology (GO) and Kyoto Encyclopedia of Genes and Genomes (KEGG) enrichment analyses were conducted using the R package clusterProfiler v4.8.3 (Wu et al., 2021). The annotations of the genes including associated GO and KEGG terms were collected from various sources: The Rice Annotation Project (RAP)(Kawahara et al., 2013) and EnsemblPlants (Martin et al., 2023) annotations for the Os-Nipponbare-Reference-IRGSP-1.0.58 reference genome, Oryzabase (Kurata and Yamazaki, 2006), the AM gene list from Das et al., 2022 and KEGG (Kanehisa and Goto, 2000; Kanehisa, 2019; Kanehisa et al., 2023). For motif enrichment analysis, promoter regions of unique gene sets were defined as 1,000 bp upstream to 100 bp downstream of each transcription start site (strand-aware); coordinates were exported in BED format, then fed to HOMER (findMotifsGenome.pl) for both de novo motif discovery (motif lengths 8, 10, and 12 bp; genomic background) and known-motif enrichment against HOMER’s built-in known motif collection (including motifs curated from public databases such as JASPAR). Enrichment statistics were obtained from HOMER outputs (knownResults.txt), with downstream visualisation and cross-comparison across gene sets performed in R, using motifs passing nominal significance thresholds (p < 0.05).

### Statistical analyses

Statistical significance of the differences between genotypes, treatments, or conditions was assessed by the parametric ANOVA test or the non-parametric Kruskal-Wallis (if the assumptions of ANOVA were not met by the dataset), at 5% significance level. Student’s T-test or Pairwise Wilcoxon rank sum were then respectively used to identify statistically significant differences between genotypes, denoted in graphs with asterisks (* for *p*-value <0.05, ** for < 0.01, *** for < 0.001 and **** for <0.0001). Validity of these tests was confirmed by a one-way analysis of variance (ANOVA) test for the log10 transformed data (to ensure equal variance). For all tests, the null hypothesis was rejected with a *p*-value threshold of 0.05. Every graph displays all data points. The exact statistical test used for each dataset is indicated in the corresponding figure legend. All statistical analyses were conducted using R (https://cran.r-project.org).

## Author contributions

R.H. and U.P. designed the study, R.H., E.W. and U.P. obtained funding; R.H., S.B., M.S.H. and E.W. generated rice transgenic material; R.H. and D.R. characterised rice transgenic material; G.F.G., R.H., D.R. performed experiments; G.F.G. performed RNA-seq data analysis, G.F.G., R.H. and D.R. analysed the data and generated figures; G.F.G., R.H. and U.P. wrote the manuscript.

## Data availability

Raw sequences, images and gene count matrices generated in this study are available in the NCBI GEO database under the accession GSE329295. R code employed to analyse the data and generate graphics is available in the following github repository: https://github.com/gabriel-ferreras/gofCCaMK_d14l

Other data supporting the findings of this article are available in the supplemental information.

## Acknowledgements

We would like to thank colleagues at the Crop Science Centre, University of Cambridge for assistance with experimental work and constructive feedback, in particular Chai Hao Chiu for valuable discussions and help shaping the project and Edwin Jarratt-Barnham for assistance in data analysis. Research was supported by the Biotechnology and Biological Sciences Research Council to E.W., R.H, G.F.G. and U.P., St John’s College (University of Cambridge) through the Benefactor’s Scholarship to G.F.G and Research Reimbursement Schemes to U.P. and G.F.G., Cambridge Philosophical Society through Research Studentship to G.F.G.

## References

Alexander T, Meier R, Toth R, Weber HC (1988) Dynamics of arbuscule development and degeneration in mycorrhizas of Triticum aestivum L. and Avena sativa L. with reference to Zea mays L. New Phytologist. doi: 10.1111/j.1469-8137.1988.tb00273.x

An J, Fang L, Cremers W, Aleksejeva K, Wang Y, Li G, Zhang M, Huang J, Ma X, Cao Q, et al (2025) A mobile DELLA controls Medicago truncatula root cortex patterning to host arbuscular mycorrhizal fungi. Nat Plants. doi: 10.1038/s41477-025-02114-6

Ané JM, Kiss GB, Riely BK, Penmetsa RV, Oldroyd GED, Ayax C, Lévy J, Debellé F, Baek JM, Kalo P, et al (2004) Medicago truncatula DMI1 Required for Bacterial and Fungal Symbioses in Legumes. Science (1979). doi: 10.1126/science.1092986

Bécard G, Fortin JA (1988) Early events of vesicular-arbuscular mycorrhiza formation on Ri T-DNA transformed roots. New Phytologist 108: 211–218

Bravo A, York T, Pumplin N, Mueller LA, Harrison MJ (2016) Genes conserved for arbuscular mycorrhizal symbiosis identified through phylogenomics. Nat Plants. doi: 10.1038/NPLANTS.2015.208

Bray NL, Pimentel H, Melsted P, Pachter L (2016) Near-optimal probabilistic RNA-seq quantification. Nat Biotechnol 34: 525–527

Cargill RIM, Shimizu TS, Kiers ET, Kokkoris V (2025) Cellular anatomy of arbuscular mycorrhizal fungi. Current Biology. doi: 10.1016/j.cub.2025.03.053

Chiu CH, Choi J, Paszkowski U (2018) Independent signalling cues underpin arbuscular mycorrhizal symbiosis and large lateral root induction in rice. New Phytologist 217: 552–557

Chiu CH, Roszak P, Orvošová M, Paszkowski U (2022) Arbuscular mycorrhizal fungi induce lateral root development in angiosperms via a conserved set of MAMP receptors. Current Biology 32: 4428–4437.e3

Choi J, Lee T, Cho J, Servante EK, Pucker B, Summers W, Bowden S, Rahimi M, An K, An G, et al (2020) The negative regulator SMAX1 controls mycorrhizal symbiosis and strigolactone biosynthesis in rice. Nat Commun. doi: 10.1038/s41467-020-16021-1

Das D, Paries M, Hobecker K, Gigl M, Dawid C, Lam H-M, Zhang J, Chen M, Gutjahr C (2022) PHOSPHATE STARVATION RESPONSE transcription factors enable arbuscular mycorrhiza symbiosis. Nat Commun 13: 477

Delaux PM, Xie X, Timme RE, Puech-Pages V, Dunand C, Lecompte E, Delwiche CF, Yoneyama K, Bécard G, Séjalon-Delmas N (2012) Origin of strigolactones in the green lineage. New Phytologist 195: 857–871

Floss DS, Levy JG, Lévesque-Tremblay V, Pumplin N, Harrison MJ (2013) DELLA proteins regulate arbuscule formation in arbuscular mycorrhizal symbiosis. Proc Natl Acad Sci U S A. doi: 10.1073/pnas.1308973110

Genre A, Lanfranco L, Perotto S, Bonfante P (2020) Unique and common traits in mycorrhizal symbioses. Nat Rev Microbiol 18: 649–660

Gleason C, Chaudhuri S, Yang T, Muñoz A, Poovaiah BW, Oldroyd GED (2006) Nodulation independent of rhizobia induced by a calcium-activated kinase lacking autoinhibition. Nature. doi: 10.1038/nature04812

Gutjahr C, Banba M, Croset V, An K, Miyao A, An G, Hirochika H, Imaizumi-Anraku H, Paszkowski U (2008) Arbuscular Mycorrhiza-Specific Signaling in Rice Transcends the Common Symbiosis Signaling Pathway. Plant Cell 20: 2989–3005

Gutjahr C, Gobbato E, Choi J, Riemann M, Johnston MG, Summers W, Carbonnel S, Mansfield C, Yang SY, Nadal M, et al (2015) Rice perception of symbiotic arbuscular mycorrhizal fungi requires the karrikin receptor complex. Science (1979) 350: 1521–1524

Gutjahr C, Radovanovic D, Geoffroy J, Zhang Q, Siegler H, Chiapello M, Casieri L, An K, An G, Guiderdoni E, et al (2012) The half-size ABC transporters STR1 and STR2 are indispensable for mycorrhizal arbuscule formation in rice. Plant Journal. doi: 10.1111/j.1365-313X.2011.04842.x

Hayashi T, Banba M, Shimoda Y, Kouchi H, Hayashi M, Imaizumi-Anraku H (2010) A dominant function of CCaMK in intracellular accommodation of bacterial and fungal endosymbionts. Plant Journal. doi: 10.1111/j.1365-313X.2010.04228.x

Hull R, Choi J, Paszkowski U (2021) Conditioning plants for arbuscular mycorrhizal symbiosis through DWARF14-LIKE signalling. Curr Opin Plant Biol 62: 102071

Jiang L, Liu X, Xiong G, Liu H, Chen F, Wang L, Meng X, Liu G, Yu H, Yuan Y, et al (2013) DWARF 53 acts as a repressor of strigolactone signalling in rice. Nature 504: 401–405

Jiang Y, Xie Q, Wang W, Yang J, Zhang X, Yu N, Zhou Y, Wang E (2018) Medicago AP2-Domain Transcription Factor WRI5a Is a Master Regulator of Lipid Biosynthesis and Transfer during Mycorrhizal Symbiosis. Mol Plant. doi: 10.1016/j.molp.2018.09.006

Kagiyama M, Hirano Y, Mori T, Kim SY, Kyozuka J, Seto Y, Yamaguchi S, Hakoshima T (2013) Structures of D14 and D14L in the strigolactone and karrikin signaling pathways. Genes to Cells 18: 147–160

Kameoka H, Kyozuka J (2015) Downregulation of Rice DWARF 14 LIKE Suppress Mesocotyl Elongation via a Strigolactone Independent Pathway in the Dark. Journal of Genetics and Genomics 42: 119–124

Kanehisa M (2019) Toward understanding the origin and evolution of cellular organisms. Protein Science 28: 1947–1951

Kanehisa M, Furumichi M, Sato Y, Kawashima M, Ishiguro-Watanabe M (2023) KEGG for taxonomy-based analysis of pathways and genomes. Nucleic Acids Res 51: D587–D592

Kanehisa M, Goto S (2000) KEGG: Kyoto Encyclopedia of Genes and Genomes.

Kawahara Y, de la Bastide M, Hamilton JP, Kanamori H, Mccombie WR, Ouyang S, Schwartz DC, Tanaka T, Wu J, Zhou S, et al (2013) Improvement of the oryza sativa nipponbare reference genome using next generation sequence and optical map data. Rice 6: 3–10

Khosla A, Morffy N, Li Q, Faure L, Chang SH, Yao J, Zheng J, Cai ML, Stanga J, Flematti GR, et al (2020) Structure–Function analysis of SMAX1 reveals domains that mediate its karrikin-induced proteolysis and interaction with the receptor KAI2. Plant Cell 32: 2639–2659

Kiers ET, Duhamel M, Beesetty Y, Mensah JA, Franken O, Verbruggen E, Fellbaum CR, Kowalchuk GA, Hart MM, Bago A, et al (2011) Reciprocal rewards stabilize cooperation in the mycorrhizal symbiosis. Science (1979) 333: 880–882

Kobae Y, Ohmori Y, Saito C, Yano K, Ohtomo R, Fujiwara T (2016) Phosphate treatment strongly inhibits new arbuscule development but not the maintenance of arbuscule in mycorrhizal rice roots. Plant Physiol 171: 566–579

Koide RT, Mosse B (2004) A history of research on arbuscular mycorrhiza. Mycorrhiza. doi: 10.1007/s00572-004-0307-4

Kurata N, Yamazaki Y (2006) Oryzabase. An integrated biological and genome information database for rice. Plant Physiol 140: 12–17

Leng J, Wei X, Jin X, Wang L, Fan K, Zou K, Zheng Z, Saridis G, Zhao N, Zhou D, et al (2023) ARBUSCULAR MYCORRHIZA-INDUCED KINASES AMK8 and AMK24 associate with the receptor-like kinase KINASE3 to regulate arbuscular mycorrhizal symbiosis in Lotus japonicus. Plant Cell. doi: 10.1093/plcell/koad050

Li XR, Sun J, Albinsky D, Zarrabian D, Hull R, Lee T, Jarratt-Barnham E, Chiu CH, Jacobsen A, Soumpourou E, et al (2022) Nutrient regulation of lipochitooligosaccharide recognition in plants via NSP1 and NSP2. Nat Commun. doi: 10.1038/s41467-022-33908-3

Liu G, Stirnemann M, Gübeli C, Egloff S, Courty PE, Aubry S, Vandenbussche M, Morel P, Reinhardt D, Martinoia E, et al (2019) Strigolactones Play an Important Role in Shaping Exodermal Morphology via a KAI2-Dependent Pathway. iScience 17: 144–154

Liu YN, Liu CC, Zhu AQ, Niu KX, Guo R, Tian L, Wu YN, Sun B, Wang B (2022) OsRAM2 Function in Lipid Biosynthesis Is Required for Arbuscular Mycorrhizal Symbiosis in Rice. Molecular Plant-Microbe Interactions. doi: 10.1094/MPMI-04-21-0097-R

Love MI, Huber W, Anders S (2014) Moderated estimation of fold change and dispersion for RNA-seq data with DESeq2. Genome Biol. doi: 10.1186/s13059-014-0550-8

Lu J, Wu T, Zhang B, Liu S, Song W, Qiao J, Ruan H (2021) Types of nuclear localization signals and mechanisms of protein import into the nucleus. Cell Communication and Signaling. doi: 10.1186/s12964-021-00741-y

Luginbuehl LH, Menard GN, Kurup S, Van Erp H, Radhakrishnan G V., Breakspear A, Oldroyd GED, Eastmond PJ (2017) Fatty acids in arbuscular mycorrhizal fungi are synthesized by the host plant. Science (1979) 356: 1175–1178

Ma H, Duan J, Ke J, He Y, Gu X, Xu TH, Yu H, Wang Y, Brunzelle JS, Jiang Y, et al (2017) A D53 repression motif induces oligomerization of TOPLESS corepressors and promotes assembly of a corepressor-nucleosome complex. Sci Adv. doi: 10.1126/sciadv.1601217

Machin DC, Hamon-Josse M, Bennett T (2020) Fellowship of the rings: a saga of strigolactones and other small signals. New Phytologist 225: 621–636

Madsen LH, Tirichine L, Jurkiewicz A, Sullivan JT, Heckmann AB, Bek AS, Ronson CW, James EK, Stougaard J (2010) The molecular network governing nodule organogenesis and infection in the model legume Lotus japonicus. Nat Commun. doi: 10.1038/ncomms1009

Martin FJ, Amode MR, Aneja A, Austine-Orimoloye O, Azov AG, Barnes I, Becker A, Bennett R, Berry A, Bhai J, et al (2023) Ensembl 2023. Nucleic Acids Res 51: D933–D941

Meng Y, Varshney K, Incze N, Badics E, Kamran M, Davies SF, Oppermann LMF, Magne K, Dalmais M, Bendahmane A, et al (2022) KARRIKIN INSENSITIVE2 regulates leaf development, root system architecture and arbuscular-mycorrhizal symbiosis in Brachypodium distachyon. Plant Journal 109: 1559–1574

Miller JB, Pratap A, Miyahara A, Zhou L, Bornemann S, Morris RJ, Oldroyd GED (2013) Calcium/calmodulin-dependent protein kinase is negatively and positively regulated by calcium, providing a mechanism for decoding calcium responses during symbiosis signaling. Plant Cell. doi: 10.1105/tpc.113.116921

Mitra RM, Gleason CA, Edwards A, Hadfield J, Downie JA, Oldroyd GED, Long SR (2004) A Ca2+/calmodulin-dependent protein kinase required for symbiotic nodule development: Gene identification by transcript-based cloning. Proc Natl Acad Sci U S A. doi: 10.1073/pnas.0400595101

Mosse B, Hayman DS, Arnold DJ (1973) Plant Growth Responses to Vesicular-Arbuscular Mycorrhiza. New Phytologist. doi: 10.1111/j.1469-8137.1973.tb02056.x

Nelson DC, Scaffidi A, Dun EA, Waters MT, Flematti GR, Dixon KW, Beveridge CA, Ghisalberti EL, Smith SM (2011) F-box protein MAX2 has dual roles in karrikin and strigolactone signaling in Arabidopsis thaliana. Proc Natl Acad Sci U S A. doi: 10.1073/pnas.1100987108

Orozco-Mosqueda MDC, Glick BR, Santoyo G (2026) Cross-talk within plant niches: endophytic and arbuscular mycorrhizal fungi for sustainable crop production. FEMS Microbiol Rev. doi: 10.1093/femsre/fuaf063

Paries M, Gutjahr C (2023) The good, the bad, and the phosphate: regulation of beneficial and detrimental plant–microbe interactions by the plant phosphate status. New Phytologist 239: 29–46

Parniske M (2008) Arbuscular mycorrhiza: The mother of plant root endosymbioses. Nat Rev Microbiol 6: 763–775

Radhakrishnan G V., Keller J, Rich MK, Vernié T, Mbadinga Mbadinga DL, Vigneron N, Cottret L, Clemente HS, Libourel C, Cheema J, et al (2020) An ancestral signalling pathway is conserved in intracellular symbioses-forming plant lineages. Nat Plants. doi: 10.1038/s41477-020-0613-7

Schindelin J, Arganda-Carreras I, Frise E, Kaynig V, Longair M, Pietzsch T, Preibisch S, Rueden C, Saalfeld S, Schmid B, et al (2012) Fiji: An open-source platform for biological-image analysis. Nat Methods 9: 676–682

Shi J, Zhao B, Zheng S, Zhang X, Wang X, Dong W, Xie Q, Wang G, Xiao Y, Chen F, et al (2021) A phosphate starvation response-centered network regulates mycorrhizal symbiosis. Cell. doi: 10.1016/j.cell.2021.09.030

Sieberer BJ, Chabaud M, Fournier J, Timmers ACJ, Barker DG (2012) A switch in Ca 2+ spiking signature is concomitant with endosymbiotic microbe entry into cortical root cells of Medicago truncatula. Plant Journal 69: 822–830

Singh S, Katzer K, Lambert J, Cerri M, Parniske M (2014) CYCLOPS, A DNA-binding transcriptional activator, orchestrates symbiotic root nodule development. Cell Host Microbe. doi: 10.1016/j.chom.2014.01.011

Smith SE, Read DJ (2008) Mycorrhizal symbiosis, Third edition edn. Academic, New York.

Soundappan I, Bennett T, Morffy N, Liang Y, Stanga JP, Abbas A, Leyser O, Nelsona DC (2015) SMAX1-LIKE/D53 family members enable distinct MAX2-dependent responses to strigolactones and karrikins in arabidopsis. Plant Cell 27: 3143–3159

Stanga JP, Morffy N, Nelson DC (2016) Functional redundancy in the control of seedling growth by the karrikin signaling pathway. Planta. doi: 10.1007/s00425-015-2458-2

Stanga JP, Smith SM, Briggs WR, Nelson DC (2013) SUPPRESSOR OF MORE AXILLARY GROWTH2 1 controls seed germination and seedling development in Arabidopsis. Plant Physiol. doi: 10.1104/pp.113.221259

Swainsbury DJK, Zhou L, Oldroyd GED, Bornemann S (2012) Calcium ion binding properties of medicago truncatula calcium/calmodulin-dependent protein kinase. Biochemistry. doi: 10.1021/bi300826m

Takeda N, Handa Y, Tsuzuki S, Kojima M, Sakakibara H, Kawaguchi M (2015) Gibberellins interfere with symbiosis signaling and gene expression and alter colonization by Arbuscular Mycorrhizal fungi in Lotus Japonicus. Plant Physiol. doi: 10.1104/pp.114.247700

Tang J, Chu C (2020) Strigolactone Signaling: Repressor Proteins Are Transcription Factors. Trends Plant Sci 25: 960–963

Tirichine L, Imaizumi-Anraku H, Yoshida S, Murakami Y, Madsen LH, Miwa H, Nakagawa T, Sandal N, Albrektsen AS, Kawaguchi M, et al (2006) Deregulation of a Ca2+/calmodulin-dependent kinase leads to spontaneous nodule development. Nature. doi: 10.1038/nature04862

Wang L, Wang B, Jiang L, Liu X, Li X, Lu Z, Meng X, Wang Y, Smith SM, Li J (2015) Strigolactone Signaling in Arabidopsis Regulates Shoot Development by Targeting D53-Like SMXL Repressor Proteins for Ubiquitination and Degradation. Plant Cell 27: 3128–3142

Wang L, Wang B, Yu H, Guo H, Lin T, Kou L, Wang A, Shao N, Ma H, Xiong G, et al (2020) Transcriptional regulation of strigolactone signalling in Arabidopsis. Nature 583: 277–281

Wang W, Shi J, Xie Q, Jiang Y, Yu N, Wang E (2017) Nutrient Exchange and Regulation in Arbuscular Mycorrhizal Symbiosis. Mol Plant 10: 1147–1158

Waters MT, Nelson DC (2023) Karrikin perception and signalling. New Phytologist 237: 1525–1541

Whiteside MD, Werner GDA, Caldas VEA, van’t Padje A, Dupin SE, Elbers B, Bakker M, Wyatt GAK, Klein M, Hink MA, et al (2019) Mycorrhizal Fungi Respond to Resource Inequality by Moving Phosphorus from Rich to Poor Patches across Networks. Current Biology 29: 2043–2050.e8

Wu T, Hu E, Xu S, Chen M, Guo P, Dai Z, Feng T, Zhou L, Tang W, Zhan L, et al (2021) clusterProfiler 4.0: A universal enrichment tool for interpreting omics data. Innovation. doi: 10.1016/j.xinn.2021.100141

Xue L, Klinnawee L, Zhou Y, Saridis G, Vijayakumar V, Brands M, Dörmann P, Gigolashvili T, Turck F, Bucher M (2018) AP2 transcription factor CBX1 with a specific function in symbiotic exchange of nutrients in mycorrhizal Lotus japonicus. Proc Natl Acad Sci U S A. doi: 10.1073/pnas.1812275115

Yang SY, Grønlund M, Jakobsen I, Grotemeyer MS, Rentsch D, Miyao A, Hirochika H, Kumar CS, Sundaresan V, Salamin N, et al (2012) Nonredundant regulation of rice arbuscular mycorrhizal symbiosis by two members of the PHOSPHATE TRANSPORTER1 gene family. Plant Cell 24: 4236–4251

Yano K, Yoshida S, Mü Ller B, J, Singh S, Banba M, Vickers K, Markmann K, White C, Schuller B, Sato S, et al (2008) CYCLOPS, a mediator of symbiotic intracellular accommodation. PNAS 105: 20540–20545

Yoshida S, Kameoka H, Tempo M, Akiyama K, Umehara M, Yamaguchi S, Hayashi H, Kyozuka J, Shirasu K (2012) The D3 F-box protein is a key component in host strigolactone responses essential for arbuscular mycorrhizal symbiosis. New Phytologist 196: 1208–1216

Zhang Q, Wang S, Xie Q, Xia Y, Lu L, Wang M, Wang G, Long S, Cai Y, Xu L, et al (2023) Control of arbuscule development by a transcriptional negative feedback loop in Medicago. Nat Commun. doi: 10.1038/s41467-023-41493-2

Zhang Y, Feng H, Druzhinina IS, Xie X, Wang E, Martin F, Yuan Z (2024) Phosphorus/nitrogen sensing and signaling in diverse root–fungus symbioses. Trends Microbiol 32: 200–215

Zheng C, Ji B, Zhang J, Zhang F, Bever JD (2015) Shading decreases plant carbon preferential allocation towards the most beneficial mycorrhizal mutualist. New Phytologist 205: 361–368

Zheng J, Hong K, Zeng L, Wang L, Kang S, Qu M, Dai J, Zou L, Zhu L, Tang Z, et al (2020) Karrikin signaling acts parallel to and additively with strigolactone signaling to regulate rice mesocotyl elongation in darkness. Plant Cell 32: 2780–2805

