## Supplemental figures for "Gain-of-function CCaMK in rice overrides genetic and anatomical barriers to arbuscular mycorrhizal colonisation"

### Supplementary figure and figure legends

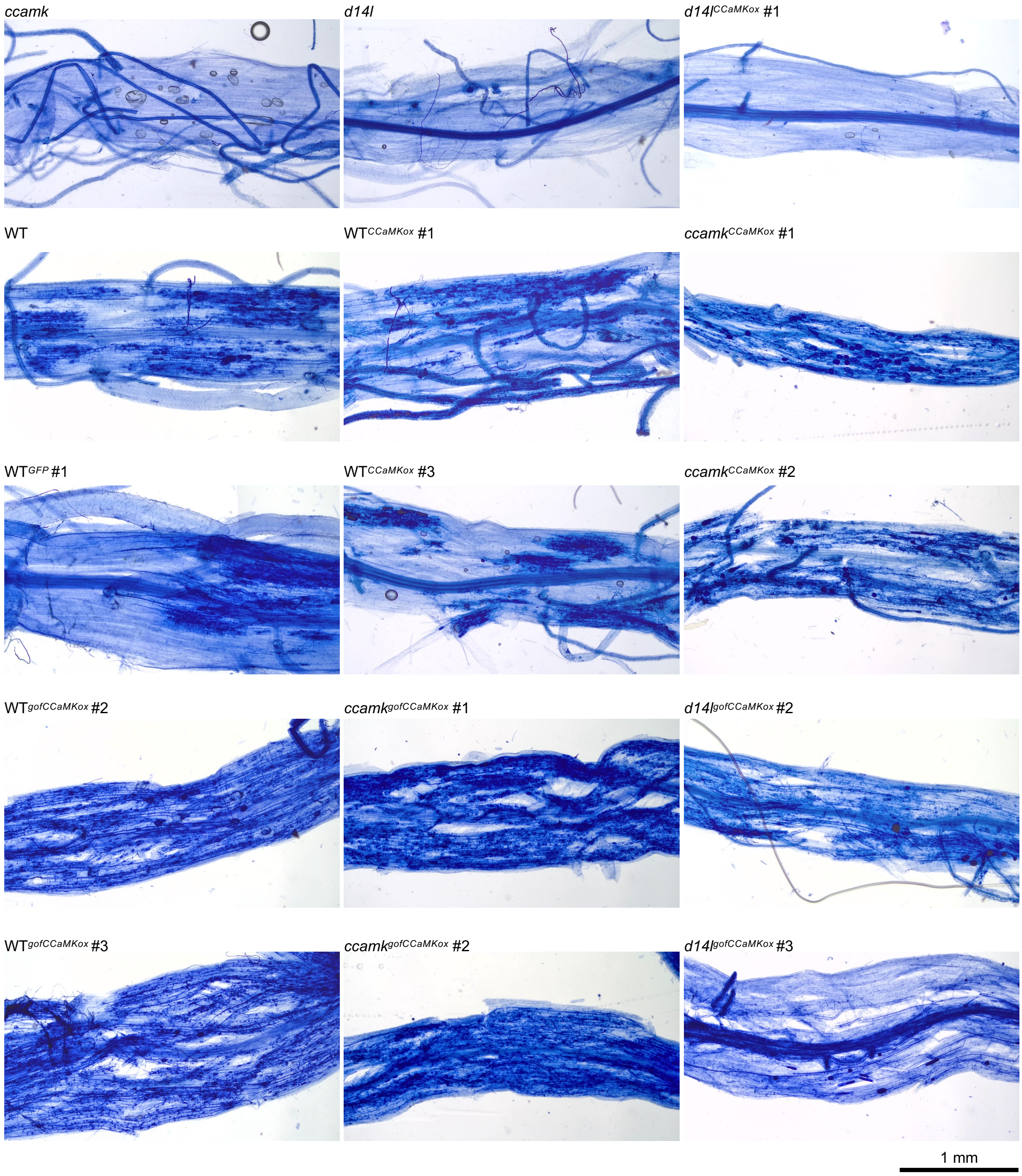


**Figure S1. Trypan-blue stained roots of *CCaMKox* and *gofCCaMKox* lines at 7 weeks-post-inoculation.** Scale bar, 1 mm.

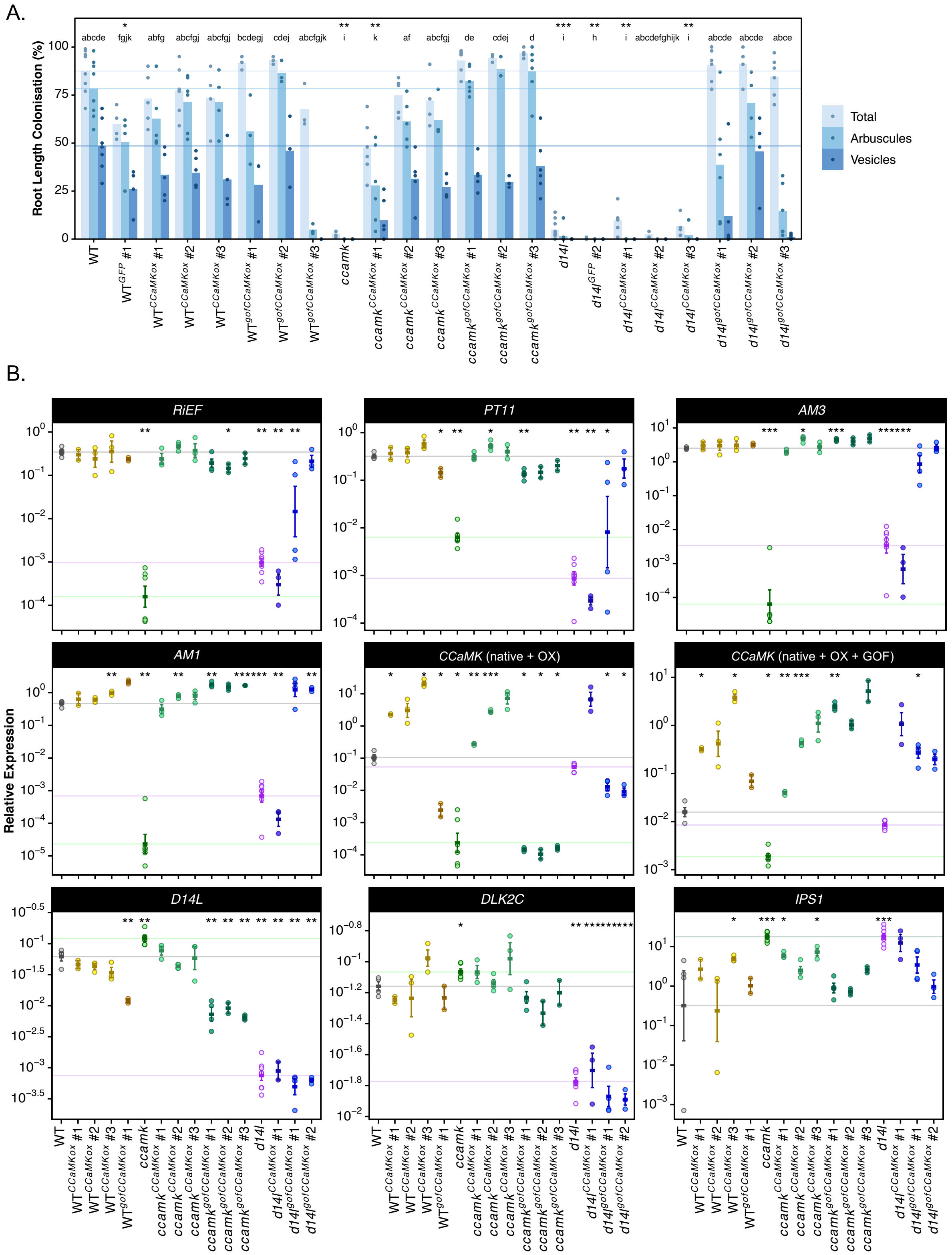


**Figure S2. Independent experiment of AM colonisation of *CCaMKox* and *gofCCaMKox* lines at 7 weeks-post-inoculation.** (A) Quantification of AM colonisation for *CCaMKox* and *gofCCaMKox* lines colonised by *R. irregularis* at 7 weeks-post-inoculation (wpi). Individual data points displayed, bars represent means for each genotype and structure. Horizontal lines show the mean for total colonisation, arbuscules and vesicles for WT. Statistically significant differences are determined by Kruskal-Wallis test followed by Pairwise Wilcoxon rank sum test. Letters denote statistically significant differences between groups ($p < 0.05$), asterisks indicate significant differences to WT (*, $p < 0.05$; **, $p < 0.01$; ***, $p < 0.001$; ****, $p < 0.0001$), regarding total colonisation. (B) Normalised gene expression levels determined via qRT-PCR for *CCaMKox* and *gofCCaMKox* lines at 7 wpi. Each graph represents gene expression of a particular gene or product defined by a primer pair (table S2), gene name or product on top, normalised to the geometric mean of three housekeeping genes. Individual datapoints are shown, error bars represent mean ± standard error. Point colour represents genotype. Horizontal lines show the mean for WT (grey), *ccamk* (green) and *d14l* (purple). Statistically significant differences are determined by Anova, followed by two-sided t-test (*, $p < 0.05$; **, $p < 0.01$; ***, $p < 0.001$; ****, $p < 0.0001$), only comparisons with WT are shown.

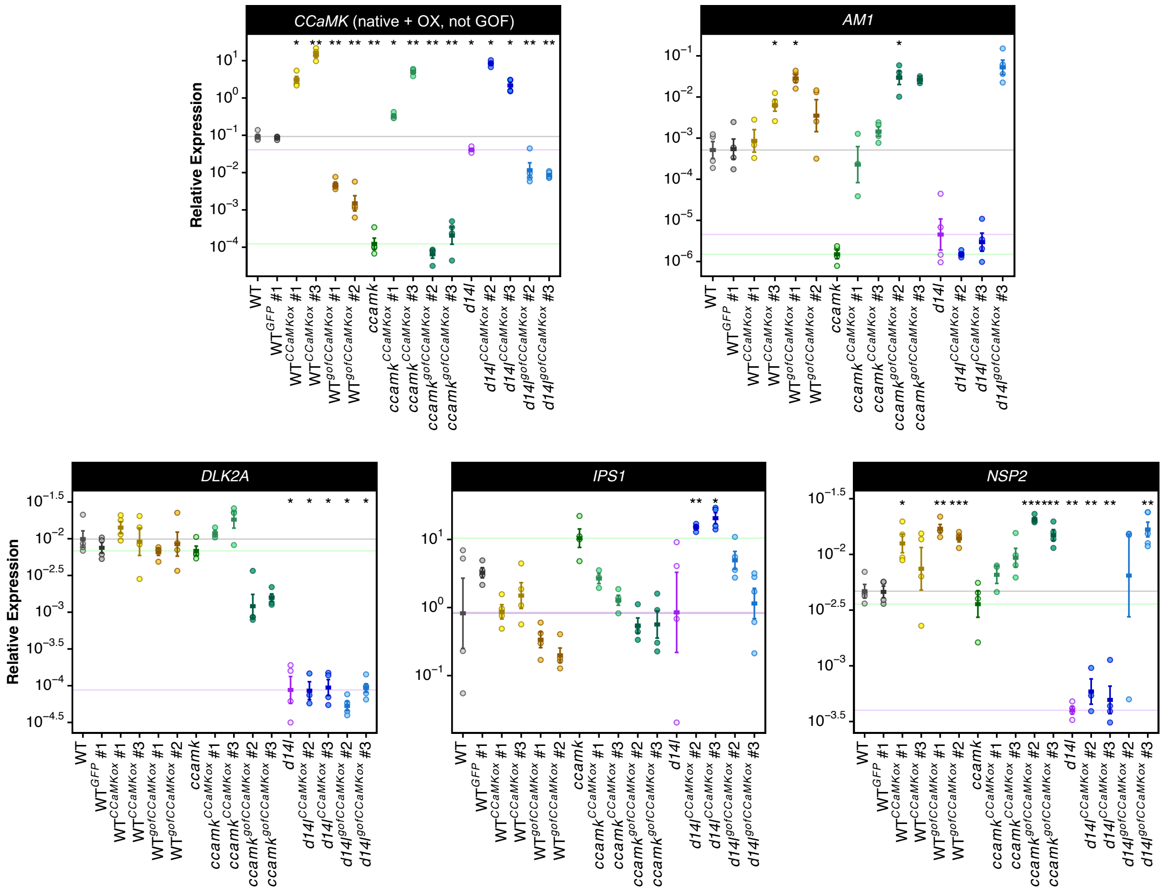


**Figure S3. Normalised gene expression levels determined via qRT-PCR for CCaMKox and gofCCaMKox lines at 7 weeks-post-inoculation for supplemental genes**. Each graph represents gene expression of a particular gene or product defined by a primer pair (table S2), gene name or product on top, normalised to the geometric mean of three housekeeping genes. Individual datapoints are shown, error bars represent mean ± standard error. Point colour represents genotype. Horizontal lines show the mean for WT (grey), *ccamk* (green) and *d14l* (purple). Statistically significant differences are determined by Anova, followed by two-sided t-test (*, $p < 0.05$; **, $p < 0.01$; ***, $p < 0.001$; ****, $p < 0.0001$), only comparisons with WT are shown.

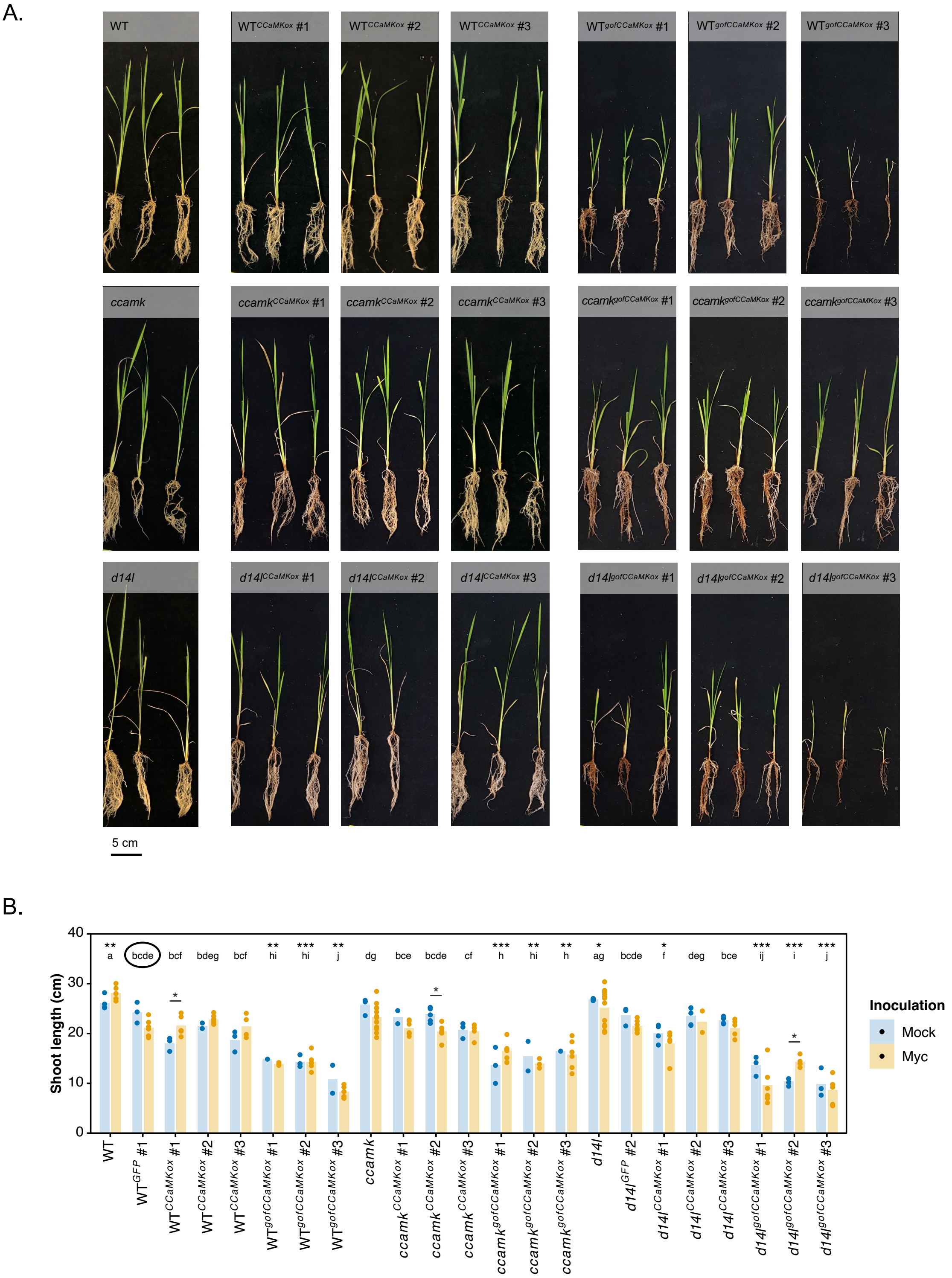


**Figure S4. *gofCCaMKox* reduces early vegetative growth in rice.** (A) Images of *CCaMKox* and *gofCCaMKox* plants at 7 weeks-post-inoculation (wpi). Plants shown were inoculated with *R. irregularis* (mock plants showed no significant differences in size). Scale bar, 5 cm. (B) Shoot length quantification for *CCaMKox* and *gofCCaMKox* lines at 7 wpi. Bars represent mean, individual data points shown. Statistically significant differences are determined by ANOVA followed by one-sided t-test. Letters denote statistically significant differences between genotype regardless of inoculation ($p < 0.05$), asterisks on top of letters indicate significant differences to $WT^{GFP}$, asterisks between bars denote significant differences between inoculated and non-inoculated (*, $p < 0.05$; **, $p < 0.01$; ***, $p < 0.001$; ****, $p < 0.0001$).

A.

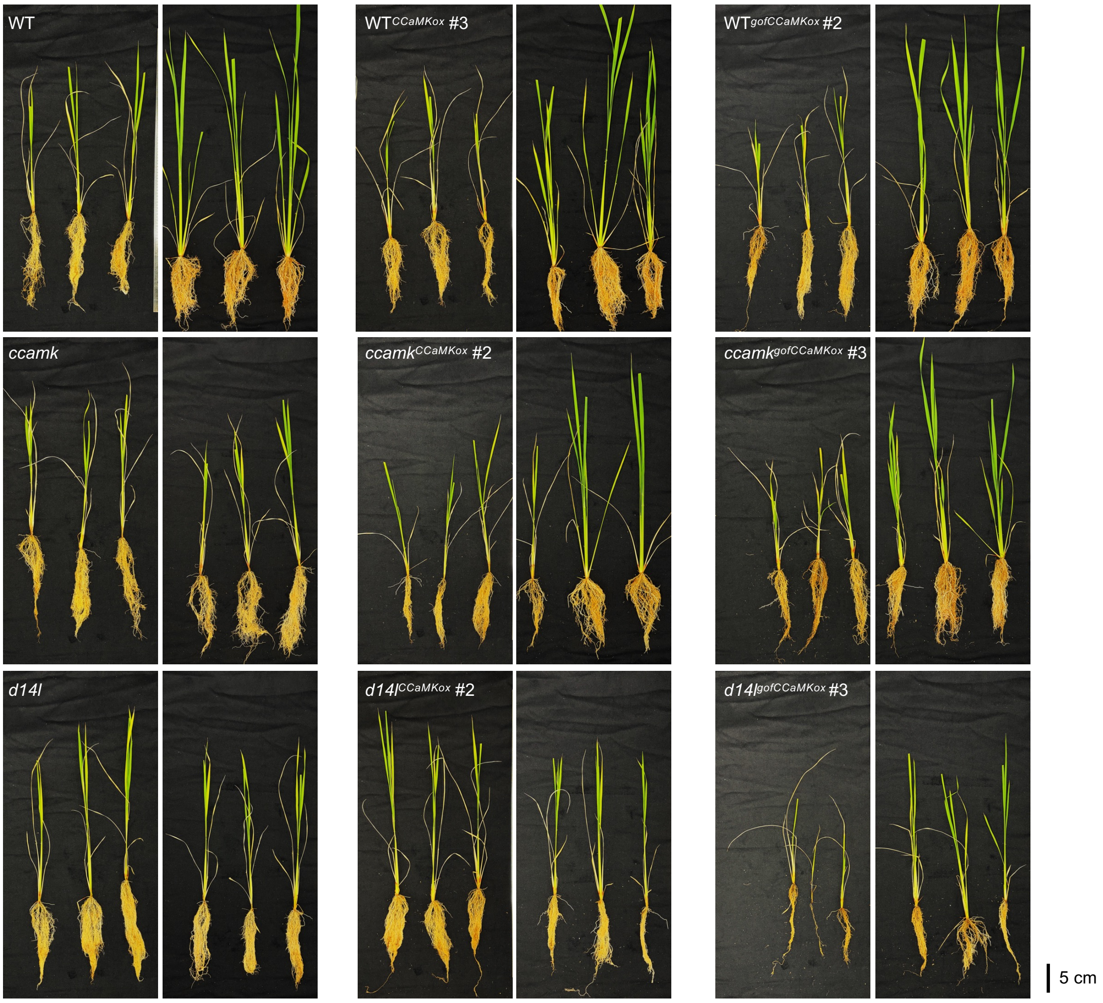


B.

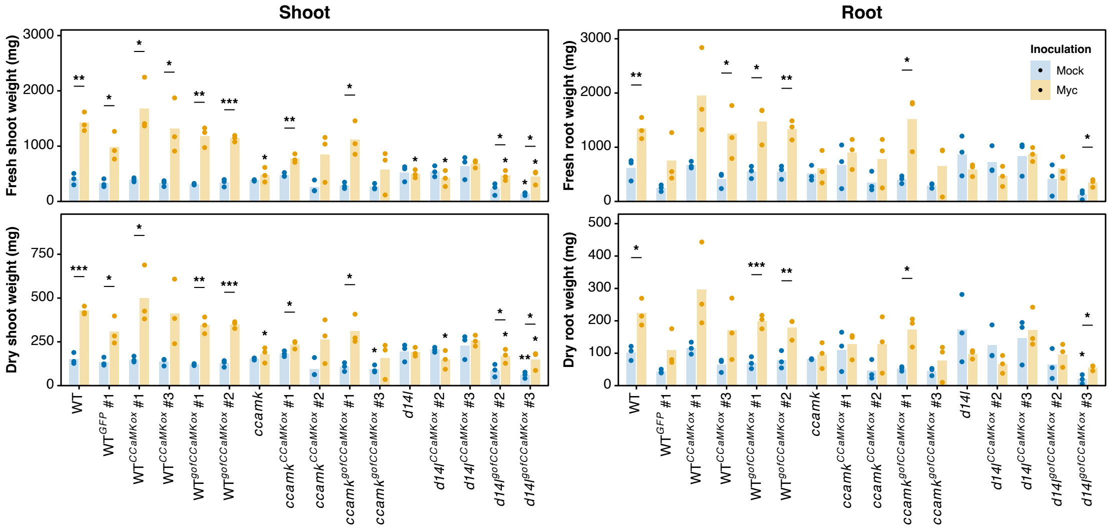


**Figure S5. *gofCCaMKox* effect on vegetative growth in rice is reduced at 12 weeks.** (A) Images of *CCaMKox* and *gofCCaMKox* plants at 12 wpi. Scale bar, 5 cm. (B) Fresh and dry shoot and root biomass quantification for *CCaMKox* and *gofCCaMKox* lines at 12 wpi. Bars represent mean ± standard error. Horizontal grey line shows the mean for $WT^{GFP}$. Statistically significant differences are determined by ANOVA followed by one-sided t-test, asterisks between bars denote significant differences between inoculated and non-inoculated, asterisks on top of bars indicate significant differences to $WT^{GFP}$ for each genotype, separately for inoculated and non-inoculated (*, $p < 0.05$; **, $p < 0.01$; ***, $p < 0.001$; ****, $p < 0.0001$).

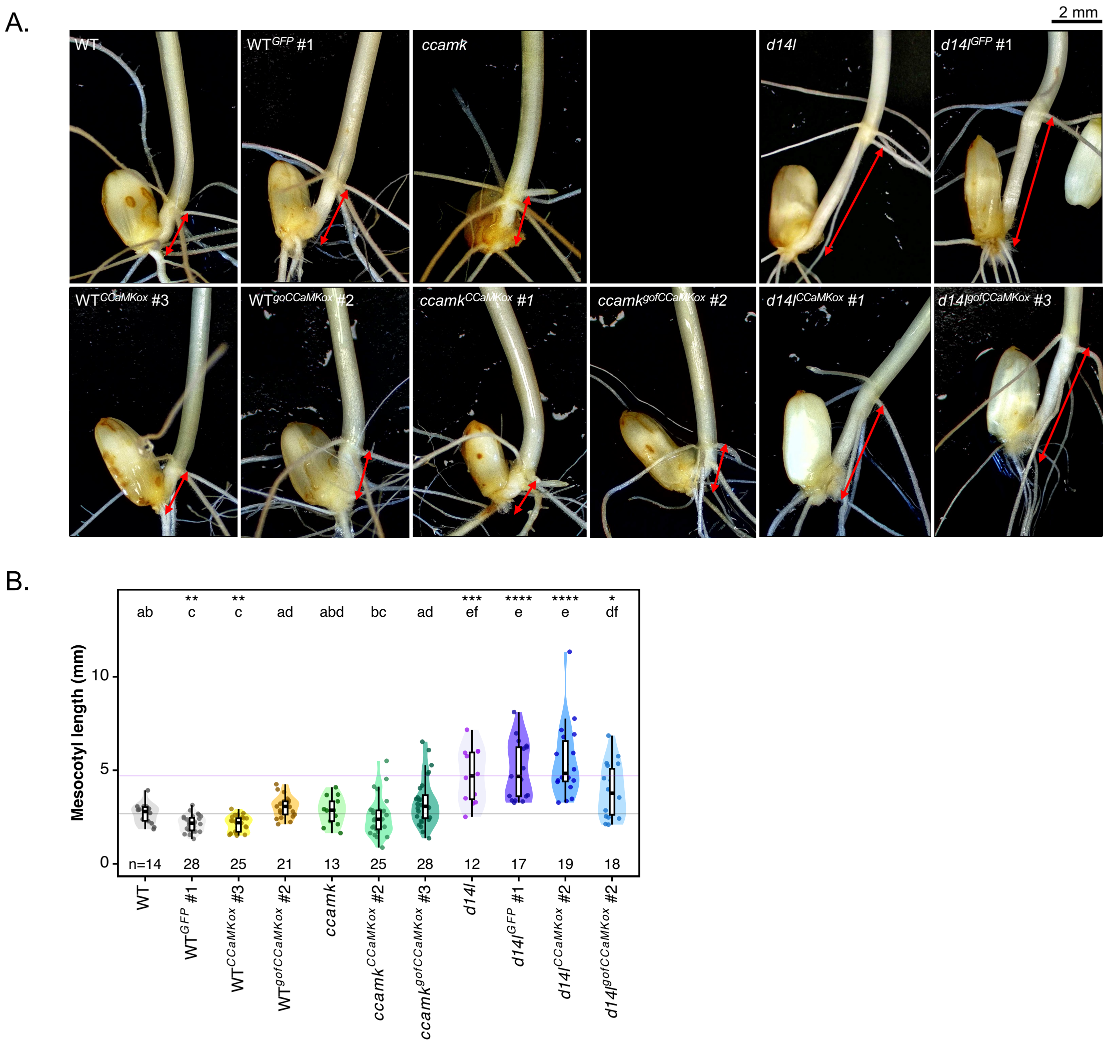


**Figure S6. Supplemental evidence for mesocotyl elongation of *CCaMKox* and *gofCCaMKox* lines at 7 days-post-imbibition.** (A) Representative stereomicroscopy images of seedlings at 7 days-post-imbibition (dpi), mesocotyl marked by red arrows. (B) Independent experiment of quantification of mesocotyl length for a subset of *CCaMKox* and *gofCCaMKox* lines at 7 dpi. Individual datapoints are shown (n > 10 seedlings, indicated on the graph). Point colour represents genotype. Horizontal lines show the mean for WT (grey) and *d14l* (purple). Statistically significant differences are determined by Kruskal-Wallis test, p-value displayed, followed by Pairwise Wilcoxon rank sum test. Letters denote statistically significant differences between groups ($p < 0.05$), asterisks indicate significant differences to WT (*, $p < 0.05$; **, $p < 0.01$; ***, $p < 0.001$; ****, $p < 0.0001$).

A.

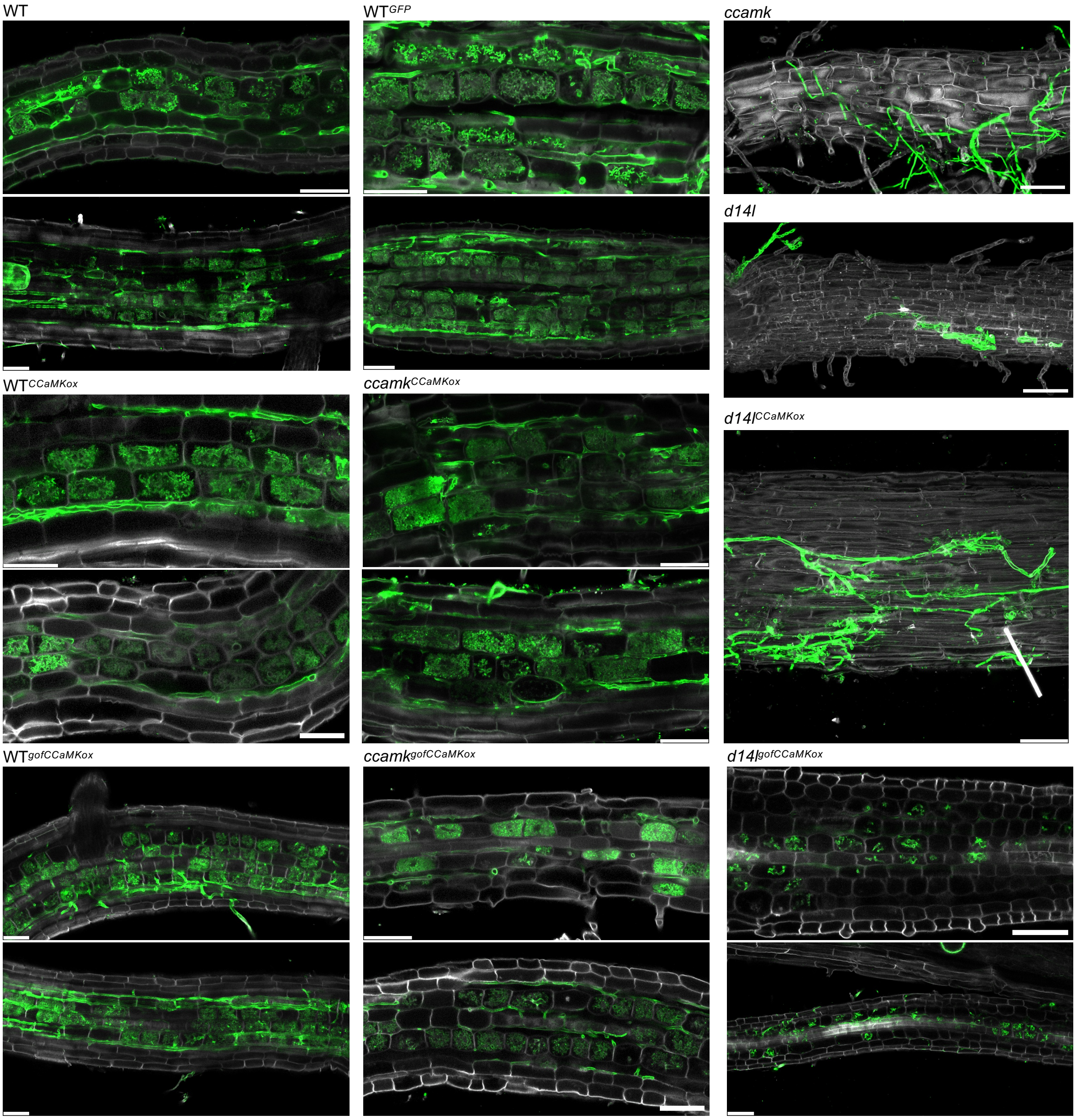


B.

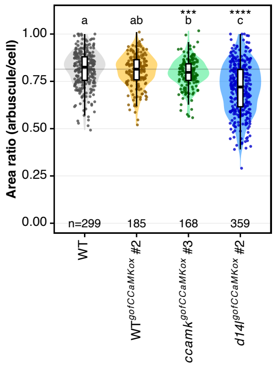


**Figure S7. Supplemental evidence for reduced arbuscule size and increased septation of intraradical hyphae in *d14l*$^{gofCCaMKox}$ roots.** (A) Supplemental CSLM images of WGA-AlexaFluor$^{488}$ stained roots of *CCaMKox* and *gofCCaMKox* lines colonised by *R. irregularis* at

7 weeks-post-inoculation (wpi) showing smaller less-branched arbuscules exclusively in *d14l*$^{gofCCaMKox}$ lines. At least two independent transformant lines, three plants, three roots and three arbuscules of each genotype were imaged. One representative image for each genotype is shown, one plane corresponding to the point of maximum arbuscule expansion. White channel corresponds to autofluorescence after excitation at 405 nm, green to WGA-AlexaFluor$^{488}$. Scale bar, 20 µm. (B) Independent experiment quantification of arbuscule size distribution of a subset of *CCaMKox* and *gofCCaMKox* lines colonised by *R. irregularis* at 7 wpi. Individual datapoints are shown ($n$ > 50 arbuscules, specified on the graph, from ≥ 3 biological replicates). Point colour represents genotype. Horizontal lines show the mean for WT (grey) and *d14l* (purple). Statistically significant differences are determined by Kruskal-Wallis test, p-value displayed, followed by Pairwise Wilcoxon rank sum test. Letters denote statistically significant differences between groups ($p$ < 0.05), asterisks indicate significant differences to WT (*, $p$ < 0.05; **, $p$ < 0.01; ***, $p$ < 0.001; ****, $p$ < 0.0001). (C)

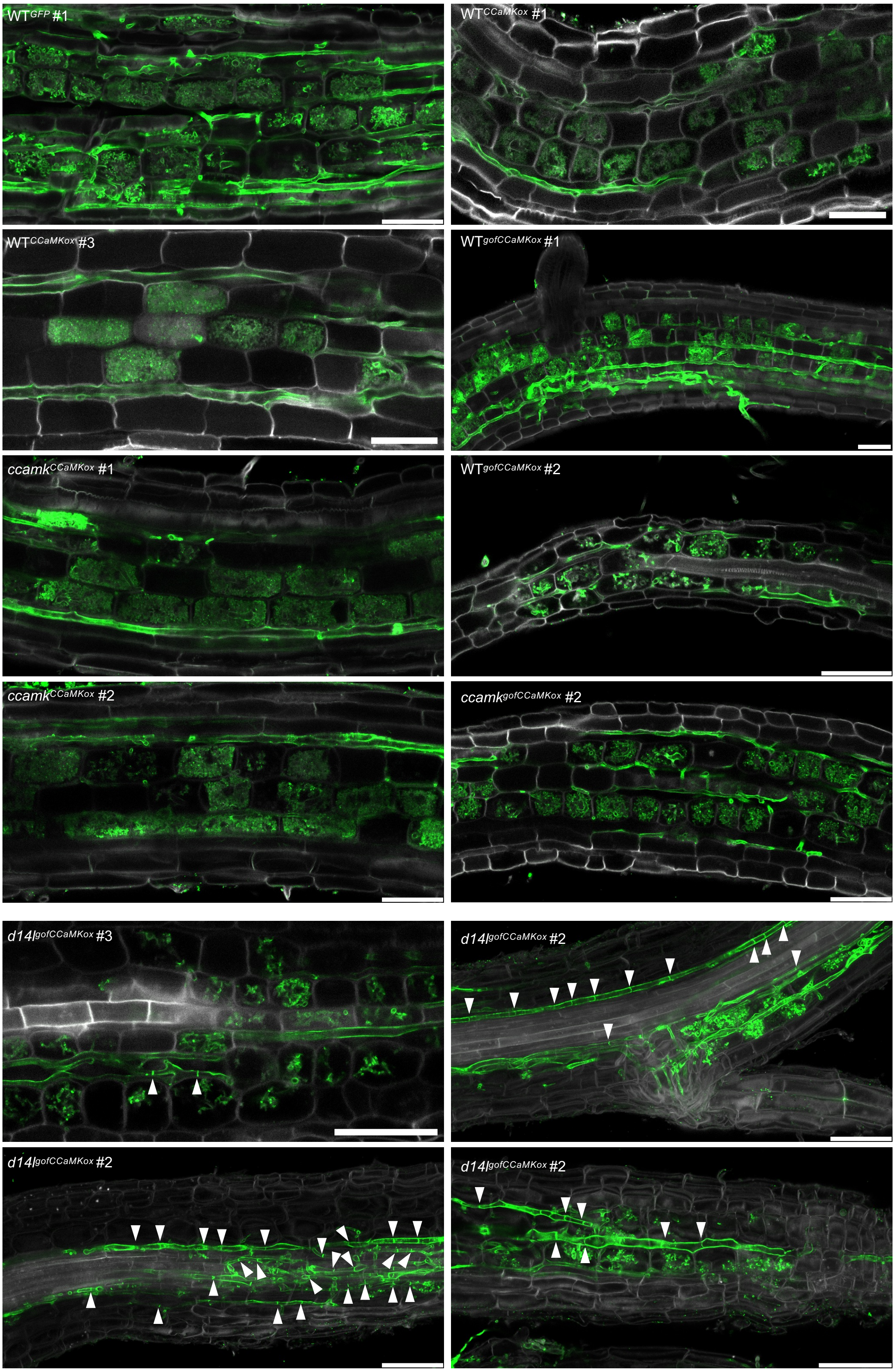


**Figure S8. Supplemental CSLM images of WGA-AlexaFluor$^{488}$ stained roots of *CCaMKox* and *gofCCaMKox* lines colonised by *R. irregularis* at 7 weeks-post-inoculation showing smaller abundant septa in intraradical hyphae within *d14l$^{gofCCaMKox}$* lines.** At least two independent transformant lines, three plants, three roots, and three arbuscules of each genotype were imaged. One representative image for each genotype of interest is shown, maximum projection of 5-10 Z planes. White channel corresponds to autofluorescence after excitation at 405 nm, green to WGA-AlexaFluor$^{488}$. White arrows highlight hyphal septa. Scale bar, 50 µm.

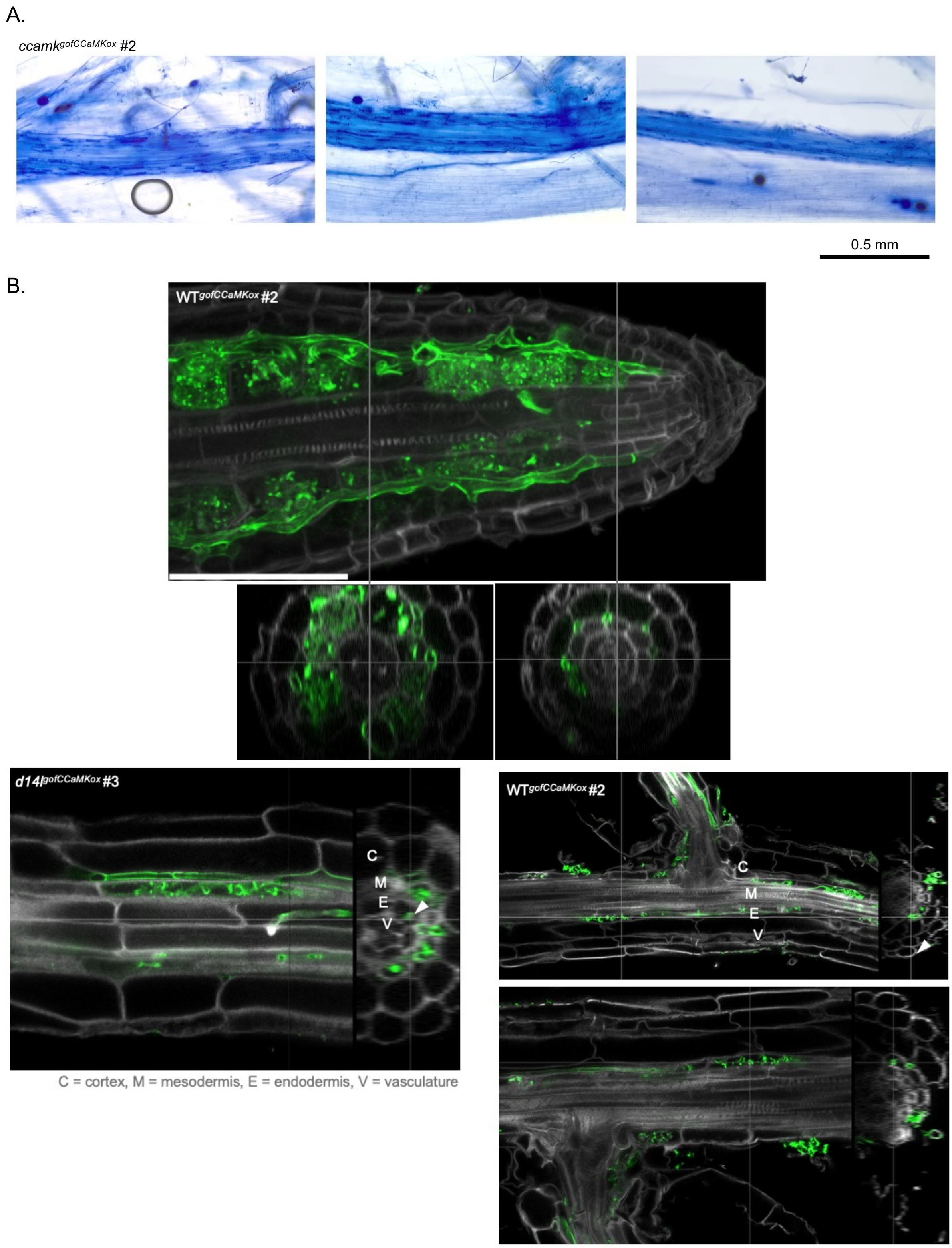


**Figure S9. Supplemental evidence that *gofCCaMKox* allows AM colonisation to expand into the root meristem and endodermis.** (A) D. Trypan-blue stained roots of *a gofCCaMKox* line at 7 weeks-post-inoculation showing arbuscules in or around the endodermis. Scale bar, 1 mm. (B) Optical reslicing of CSLM images of WGA-AlexaFluor$^{488}$ stained root tips gofCCaMKox lines colonised by *R. irregularis* at 7 weeks-post-inoculation, showing AM colonisation of the endodermis and formation of arbuscules in endodermal cells. At least two independent transformant lines, three plants, three roots, and three arbuscules of each genotype were imaged. White channel corresponds to autofluorescence after excitation at 405 nm, green to WGA-AlexaFluor$^{488}$. Scale bar, 50 µm.

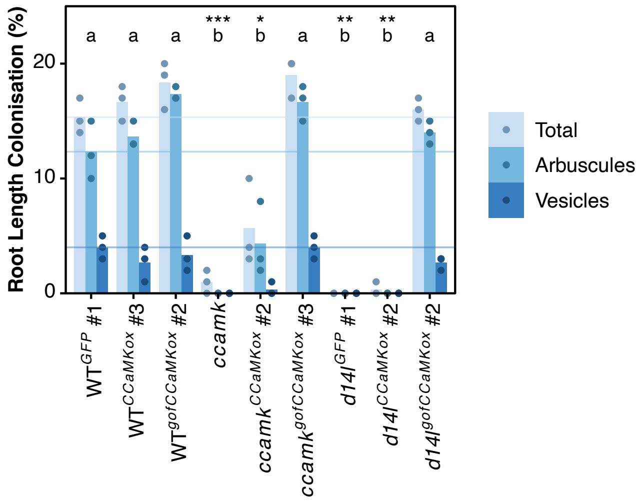


**Figure S10. Quantification of AM colonisation for *CCaMKox* and *gofCCaMKox* lines colonised by *R. irregularis* at 3 weeks-post-inoculation (wpi).** Individual data points displayed, bars represent means for each genotype and structure. Horizontal lines show the mean for total colonisation, arbuscules, and vesicles for WT. Statistically significant differences are determined by Kruskal-Wallis test followed by Pairwise Wilcoxon rank sum test. Letters denote statistically significant differences between groups ($p < 0.05$), asterisks indicate significant differences to WT (*, $p < 0.05$; **, $p < 0.01$; ***, $p < 0.001$; ****, $p < 0.0001$), regarding total colonisation.

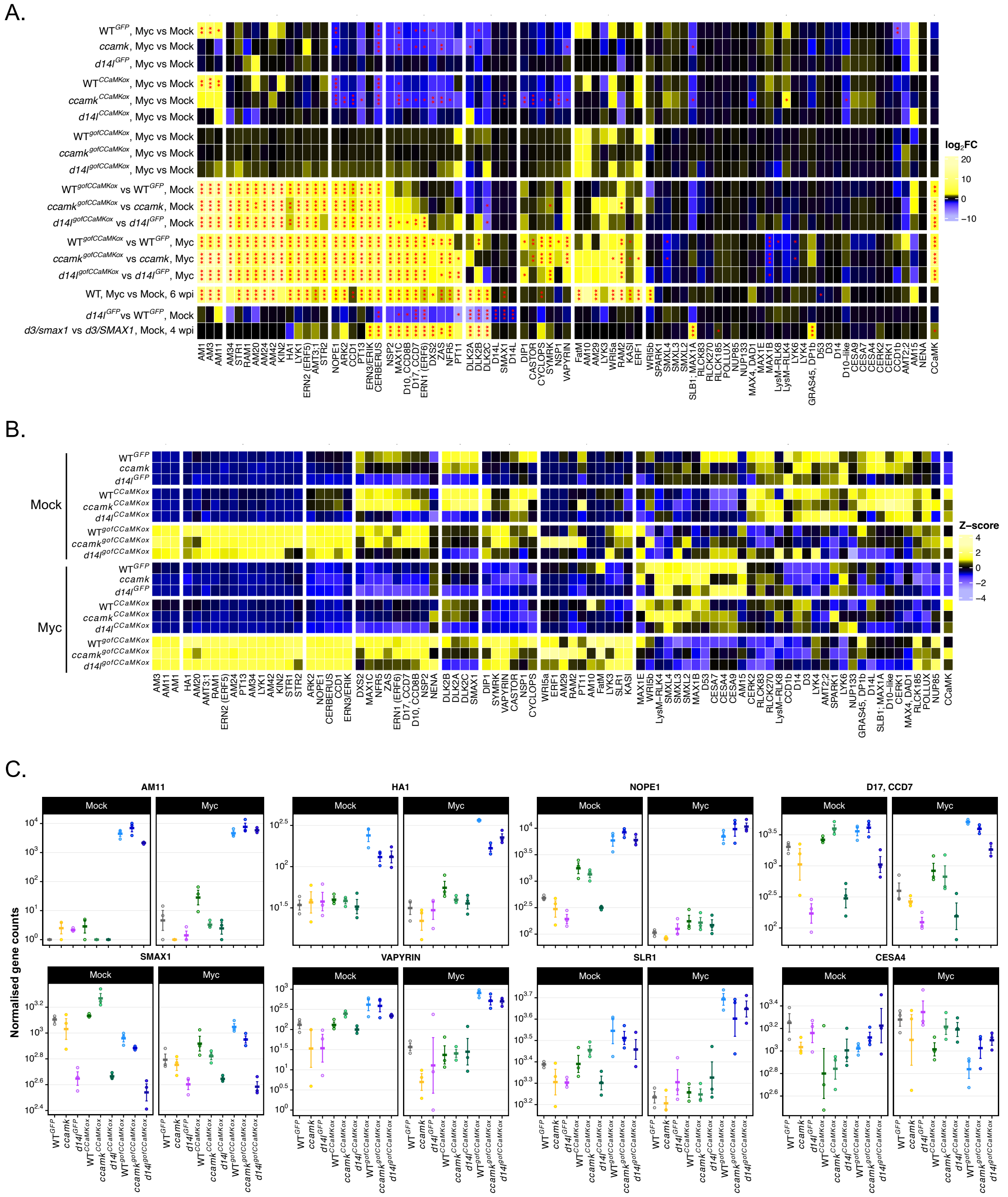

**Figure S11. Transcriptomic profiles for AM-related genes in *CCaMKox* and *gofCCaMKox* at 3 weeks-post-inoculation.** (A) Heatmap of $log_2FC$ for comparisons of interest for a selection of genes related to AM symbiosis (subset of the AM gene list from Das et al., 2022; common names shown, gene IDs in table S11). Significance levels as determined by DEG using DESeq2 shown with asterisks (* for *p*-value <0.05, ** for < 0.01, *** for < 0.001). Genes were manually curated into 8 modules based on expression profiles. (B) Heatmap of scaled VST-transformed counts for the same selection of genes related to AM symbiosis (C). DESeq2 normalised gene counts of a gene from each module.

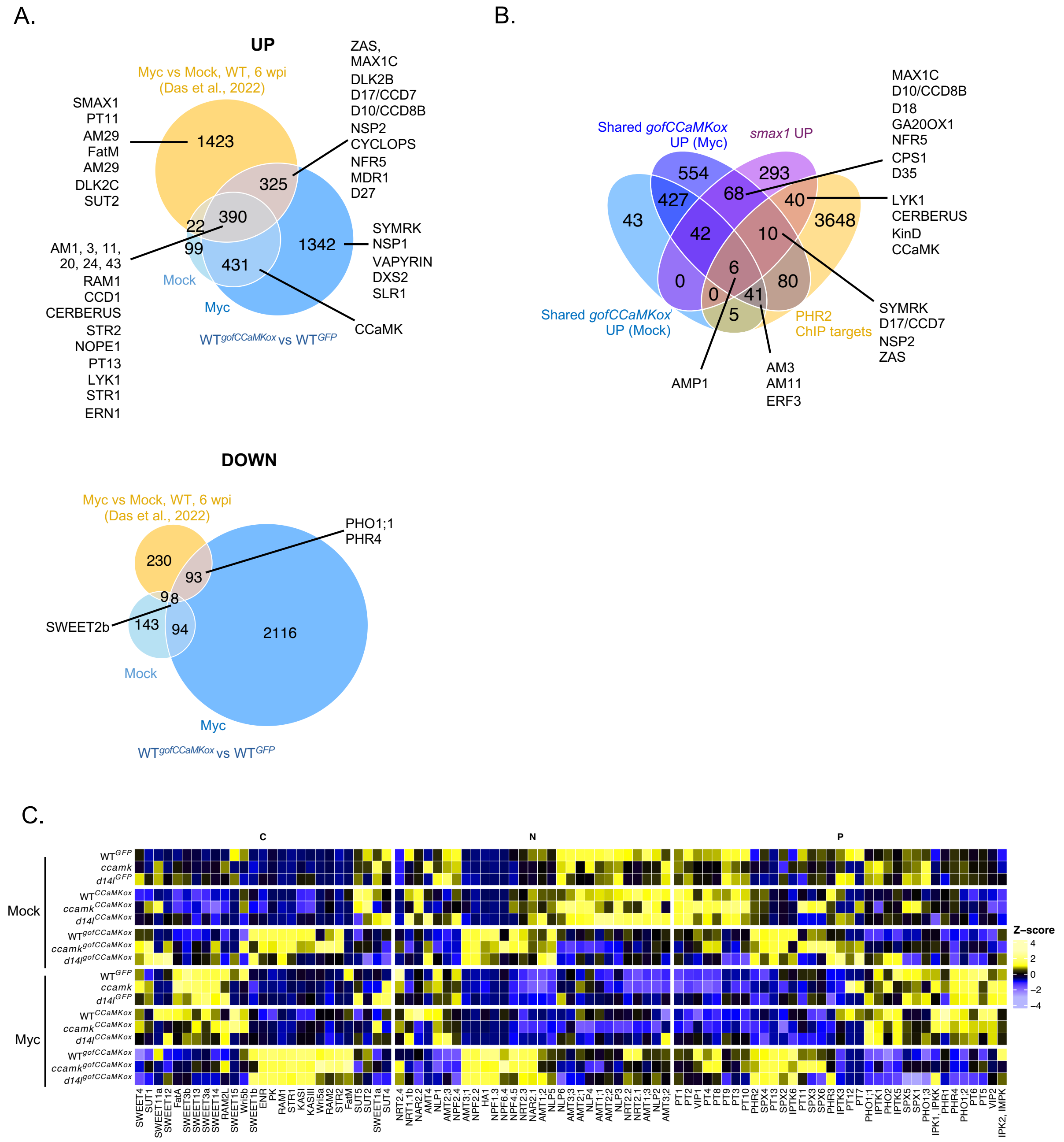


**Figure S12. gofCCaMK drives excessive gene activation and repression that does not occur even at later stages of AM colonisation.** (A) Venn diagram showing overlaps between induced and repressed genes for mycorrhizal versus non-mycorrhizal roots at 6 weeks-post-inoculation (Das et al., 2022) with DEGs for WT$^{gofCCaMKox}$ versus its background control WT$^{GFP}$, in both mock and myc conditions. Genes of interest highlighted. (B) Venn diagram showing overlaps between genes commonly up-regulated by *gofCCaMKox* in all background, either in myc or mock conditions, with the *smax1* UP gene set from Choi et al., 2020, and the PHR2 targets identified by ChIP-seq from Das et al., 2022. Genes of interest highlighted. (C) Heatmap of scaled VST-transformed counts for genes involved in nutrient transport, perception or signalling, classified in three categories based on the nutrient:

phosphorus (P), nitrogen (N) or carbon (C). Common gene names displayed, gene IDs available in table S15.

**Supplementary table legends**

**Table S1.** Transgenic rice lines generated and used in this study.

**Table S2.** Oligonucleotides and DNA sequences used in this study**.**

**Table S3.** AM colonisation scoring at 3 and 7 weeks-post-inoculation (raw data).

**Table S4.** RT–qPCR measurements (raw data).

**Table S5.** Plant height at 7 weeks-post-inoculation (raw data).

**Table S6.** Fresh and dry biomass at 12 weeks-post-inoculation (raw data).

**Table S7.** Mesocotyl length at 7 days-post-imbibition (raw data).

**Table S8.** Arbuscule abundance/ratio at 7 weeks post inoculation (raw data).

**Table S9.** Hyphal distance to the quiescent centre at 7 weeks-post-inoculation (raw data).

**Table S10.** DESeq2-normalised gene counts for all RNA-seq samples.

**Table S11.** AM-associated gene list (subset from Das et al., 2022).

**Table S12.** Differentially expressed genes in gofCCaMKox lines versus background genotypes (overlaps).

**Table S13.** Promoter motif enrichment for genes commonly induced/repressed by *gofCCaMKox*, PHR2-ChIP targets and *smax1* up and down genes.

**Table S14.** Differentially expressed genes in d14lgofCCaMKox versus WTgofCCaMKox.
